# STAG2 Maintains Chromatin Compartmentalization and Represses Regulatory Element Contact to Promote Oncogenic Signaling in Muscle Invasive Bladder Cancer

**DOI:** 10.64898/2026.06.10.731379

**Authors:** Sarah R. Athans, Jiaojiao Zhou, Xiaozhuo Liu, Eduardo Cortes-Gomez, Bhavisha Doshi, Justine Jacobi, Aimee Stablewski, Sofia Lage-Vickers, Pablo Sanchis, Maria Pia Valacco, Dean G. Tang, Geraldine Gueron, Tao Liu, Anna Woloszynska

## Abstract

Contrary to other cancer types, stromal antigen 2 (STAG2) expression is associated with shorter survival and an invasive phenotype in muscle invasive bladder cancer (MIBC). As a cohesin complex component, STAG2 regulates genome organization and cell type-specific transcription, yet its mechanistic role in MIBC remains unclear. Here, we uncover mechanisms through which STAG2 coordinates chromatin architecture and gene regulation in MIBC. Modulation of STAG2 rewired chromatin architecture, altering chromatin contacts and compartmentalization, increasing promoter-enhancer interactions, and reducing short-ranged chromatin loops. At specific gene loci, we discovered that STAG2 has context-specific activating and repressive regulatory functions. At the STAG2-activated gene *ABCA1,* STAG2 maintained A-compartment chromatin and high levels of promoter acetylation, indicators of active transcription. STAG2 KO resulted in loss of promoter acetylation, a shift from A to B compartment chromatin, and increased occupancy of the co-repressor TRIM28, resulting in *ABCA1* downregulation and diminished invasive potential. Conversely, at the STAG2-repressed gene *SPOCK3,* STAG2 KO resulted in B to A compartment switching, aberrant formation of chromatin loops, and *SPOCK3* upregulation. Treatment with EZH2 inhibitor tazemetostat augmented STAG2-KO induced *SPOCK3* upregulation, suggesting a collaborative role of STAG2 and EZH2 in repressing Polycomb Repressive Complex 2 (PRC2) target genes. Altogether, our results indicate that STAG2 plays a multifaceted role in regulating gene expression in bladder cancer that is dictated by the epigenetic and chromatin landscape of the cells. These findings identify STAG2-dependent vulnerabilities and provide a rationale for therapeutic targeting of chromatin regulators in MIBC.

## BACKGROUND

The cohesin complex member stromal antigen 2 (STAG2) is of significant interest in cancer, as one of only twelve genes frequently mutated in four or more cancer types [1]. STAG2 is considered a tumor suppressor protein in most cancer contexts, including Ewing sarcoma, acute myeloid leukemia, pancreatic ductal adenocarcinoma, and lung adenocarcinoma, as loss of function mutations in STAG2 lead to more aggressive disease and worse patient outcomes [2–7].

STAG2 is also frequently mutated in bladder cancer, and most mutations are truncating in nature leading to loss of STAG2 protein expression [8, 9]. STAG2 mutations are more frequent in lower-stage, lower-grade non-muscle invasive bladder cancer (NMIBC) compared to the aggressive muscle invasive bladder cancer (MIBC). Paradoxically and in contrast with the tumor suppressive function of STAG2 in other cancer types, STAG2 expression is associated with worse survival for muscle invasive bladder cancer (MIBC) patients and is an independent predictor of progression from non-muscle invasive bladder cancer (NMIBC) to MIBC [10, 11]. Additionally, we have previously found that STAG2 expression accelerates bladder tumor growth *in vivo* and supports an invasive phenotype *in vitro,* supporting a proposed pro-tumor role of STAG2 in MIBC [10]. This suggests that in bladder cancer, specifically, STAG2 may have a tumor promoting function. However, the mechanism by which STAG2 supports an invasive phenotype and promotes MIBC, in contrast to the tumor suppressive function of STAG2 in other cancer types, remains unknown.

In non-malignant cells, STAG2 functions as a member of the ring-shaped four-member cohesin complex, which facilitates sister chromatid cohesion prior to mitosis [12–17]. A more recently appreciated function of the cohesin complex is in topological organization of 3D genome in the interphase nucleus. Through this function, cohesin can dictate chromatin compartmentalization, maintain topologically associated domains (TADs), and form chromatin loops all of which can influence gene expression [3–6, 18–23]. TADs are genomic domains flanked by insulator regions which constrain contact between regulatory elements, such as enhancers and promoters, preventing off-target gene regulatory activity from occurring between distant regions [24]. Similarly, cohesin-mediated chromatin loops can maintain proximity of regulatory elements to promote gene expression or enforce insulation between regions to prevent aberrant gene activation [6, 19–21]. We have found that modulation of STAG2 expression does not impair the sister chromatid cohesion functions of cohesin in MIBC cells [10]. However, STAG2 knockdown (KD) is associated with alterations in gene expression, including downregulation of extracellular matrix related gene signatures, demonstrating an impact on the gene-regulatory function of cohesin [10]. Therefore, STAG2 may exert a pro-oncogenic function in MIBC through regulating gene expression, via cohesin-STAG2-facilitated chromatin organization and enforcement of regulatory element interactions throughout the genome.

STAG2 and the cohesin complex can also collaborate with epigenetic regulators, such as TRIM28, the PRC2 complex, and the PRC1 complex to exert gene regulatory functions [5, 19, 25–27]. Previous research indicates a role for cell-type-specific regulation of gene expression by STAG2 [4, 18–21]. However, the mechanisms which explain the cell-type specific localization and gene regulatory function of STAG2 remain unknown. Given the established role for cohesin in chromatin organization, we hypothesized that STAG2 promotes MIBC by maintaining chromatin architecture that supports pro-invasive gene expression programs and represses tumor-suppressive gene expression programs.

In the present study, we sought to determine how STAG2 regulates chromatin architecture, whether cooperation with epigenetic regulators contributes to the pro-invasive activity of STAG2, and how this function influences the expression of oncogenic and tumor suppressive gene expression programs specifically in MIBC. We evaluated STAG2-specific features using integrated transcriptomic, chromatin binding, proteomic, and Hi-C analyses of clinical samples and *in vitro* model systems with modulation of STAG2 expression. Our findings uncover novel characteristics of STAG2 function which may be investigated in the future as STAG2-specific therapeutic opportunities in MIBC.

## MATERIALS AND METHODS

### Roswell Park Comprehensive Cancer Center (Roswell Park) patient cohort

Tumor samples from patients with bladder cancer and with informed consent were collected at the time of radical cystectomy at Roswell Park as described previously [28].

### IPA Canonical Pathway Analysis

Differentially expressed genes (DEGs) between tumor (n = 66) and normal (n = 15) samples were identified from RNA-seq data using a threshold of false discovery rate (FDR) ≤ 0.05 and absolute fold change (FC) ≥ 4. The resulting DEG list was used for downstream pathway analysis.

Canonical pathway enrichment analysis was performed using QIAGEN Ingenuity Pathway Analysis (IPA; QIAGEN Inc., Redwood City, CA, USA) [29]. Gene identifiers and corresponding expression values (log2 fold change) were uploaded into IPA and mapped to the Ingenuity Knowledge Base. Canonical pathways significantly associated with the dataset were identified based on two metrics: (i) a right-tailed Fisher’s exact test, which evaluates the probability that the association between the dataset and a given pathway occurs by chance, and (ii) the ratio of genes from the dataset that map to a pathway relative to the total number of genes in that pathway. Pathways with p < 0.05 were considered significantly enriched and were ranked based on – log10(p-value) for visualization.

### Cell Culture

T24 cells (ATCC Cat# HTB-4) were cultured in McCoy’s medium and TCC-SUP (ATCC Cat# HTB-5) in MEM plus 1x MEM non-essential amino acid. Media for all lines was supplemented with 10% fetal bovine serum and penicillin/streptomycin. All media was purchased from Corning.

All cell lines were maintained at 5% CO_2_, 37 °C. Cells were tested for mycoplasma via Mycale Mycoplasma Detection Kit (Lonza, Cat#LT07-218) after thawing and minimally once every three months thereafter. All experiments were performed using cell lines passage 20 or below.

### CRISPR-Cas9 Knockout of STAG2

Two single-guide RNA (sgRNA) sequences were designed using the CRISPOR (tetof.net) prediction website. These guides were used to target human STAG2 exons 5 and 8 in T24 and TCCSUP cells: TCTGGTCCAAACCGAATGAA TGG (e5), GATTATCCACTTACCATGGC TGG (e8). CRISPR RNA (crRNA) and tracer RNA (trRNA) were purchased from IDT DNA Technologies (Coralville, IA) and resuspended to 160 μM each. crRNA and trRNA (1:1) were complexed using touchdown polymerase chain reaction (PCR) (IDT DNA Technologies), and Cas9 3NLS protein (61uM) (IDT DNA Technologies) was added to make functional ribonucleoprotein (RNP). RNPs for both exons 5 and 8 were added to 3 × 10^6^ cells suspended in Opti-MEM medium (ThermoFisher Scientific). The cell mixture was electroporated using a NEPA21 electroporator (Bulldog Bio, Portsmouth, NH) and then transferred into 6-well plates containing complete medium without antibiotics (2 ml). Cells were single cell sorted into 96 well plates then expanded for knockout confirmation. T24 cells were analyzed for knockout by targeted next generation sequencing and deletion analysis by Washington University at St. Louis’ Genome Editing Core facility, and a large deletion was shown between exons 5 and 8.

### Lentivirus-Mediated Knockdown of TRIM28

pAPM-D4 miR30-TRIM28 ts1 and pAPM-D4- miR30-L1221 were purchased via Addgene from Jeremy Luban (Addgene plasmid # 115862; http://n2t.net/addgene:115862; RRID: Addgene_115862 and Addgene plasmid # 115846; http://n2t.net/addgene:115846; RRID: Addgene_115846) [30]. Human embryonic kidney cells, line 293TN (System Biosciences, Cat # LV900A-1), were grown in 6 well plates Dulbecco’s modified Eagle’s medium (Invitrogen) containing 10% fetal bovine serum and 0.1% Penicillin-Streptomycin. The cells were cultured to 90-95% confluence and co-transfected with 2.5 μg of the shTRIM28/non-targeting control lentiviral constructs and 10 μg of the pPACKH1-plasmid mix (System Biosciences, Cat #LV500A-1) the TransIT-X2 Dynamic Delivery System (Mirus Bio, Product # MIR 6004). The viral supernatant was collected at both 48 h and 72 h after transfection and filtered using a 0.45μM filter. T24 cell lines (A6 Control, G2 STAG2 KO, and H2 STAG2 KO) were infected with lentivirus in the presence of 8μg/ml of polybrene (Santa Cruz Biotech, Cat # SC-134220). Transduced cells were enriched by puromycin selection for 1 week.

### Western Blot

Cells were cultured to 75-80% confluence and lysed in Triton X-100/SDS lysis buffer (1% Triton X-100, 0.1% SDS, 50mM Tris, 150mM NaCl) containing protease inhibitors (Roche). Protein concentrations were determined using Bio-Rad protein assay (Biorad, Cat #500-0116). Equal amounts of protein lysates (STAG2 IB: 50 µg; TRIM28 IB: 10 µg; ABCA1 IB: 125 µg) were prepared and mixed with 6x loading buffer. Samples were boiled for 10 minutes at 95C, except in the case of ABCA1 IB, where samples were NOT boiled. Samples were resolved on a 4-20% gradient SDS-PAGE and electro transferred onto PVDF membranes via the wet transfer method. Blots were blocked with 5%milk-TBS for at least one hour. Blots were incubated with antibodies as listed in **Table 1** overnight at 4°C.

**Table 1.** Antibodies utilized for western blotting in the current study.

| Protein (Mfg, Catalog Number) | Dilution | Diluent |
| --- | --- | --- |
| STAG2 (Cell Signaling, 5882S) | 1:500 | 5 % Milk-TBST |
| GAPDH (Abcam, ab9485) | 1:10,000 | 5 % Milk-TBST |
| TRIM28 (Proteintech, 15202-1-AP) | 1:4,000 | 5 % Milk-TBST |
| ABCA1 (Cell Signaling, 96292T) | 1:200 | 5 % Milk-TBST |
| H3K27me3 (Cell Signaling, 9733S) | 1:1,000 | 0.5 % Milk-TBST |
| Total H3 (Cell Signaling, 9715S) | 1:1,000 | 5 % Milk-TBST |
| Anti-Rabbit IR Secondary (LICOR Bio, 926-32213) | 1:5,000 | 5 % Milk-TBST |
| Anti-Mouse IR Secondary (LICOR Bio, 925-68070) | 1:4,000 | 5 % Milk-TBST |

Signal was detected using the LICOR Odyssey CLx Imager. If necessary, membranes were stripped using Restore Western Blot Stripping Buffer (Thermo Scientific, Cat #46430) then re-blocked for at least one hour prior to probing for the next protein of interest.

### RNA Isolation and cDNA Synthesis

Total RNA was extracted using the Direct-zol RNA MiniPrep kit (Zymo Research, Cat #R2052) and quantified using a Nanodrop 8000 system (Thermo Scientific). cDNA was synthesized from 0.5-1 µg RNA using the iScript cDNA synthesis kit (BioRad, Cat #170-8891).

### qPCR

10 µL qPCR reactions were prepared using PowerTrack SYBR green (Applied Biosystems, Cat No. A46109) according to manufacturers’ instructions. Primers were designed for target genes or genomic regions as noted in Table 2. 500 nM each of forward and reverse primers were added to each reaction in addition to 50 ng cDNA (for RNA analysis) or 1 µL ChIP DNA (for ChIP experiments). Samples were plated in at least technical duplicates, and the PCR reaction was run for 40-45 cycles depending on the amplicon on the CFX Connect Real-Time PCR Detection System (Biorad). Results were calculated and reported using the delta-delta-Ct method relative to the indicated control for each experiment.

**Table 2.**
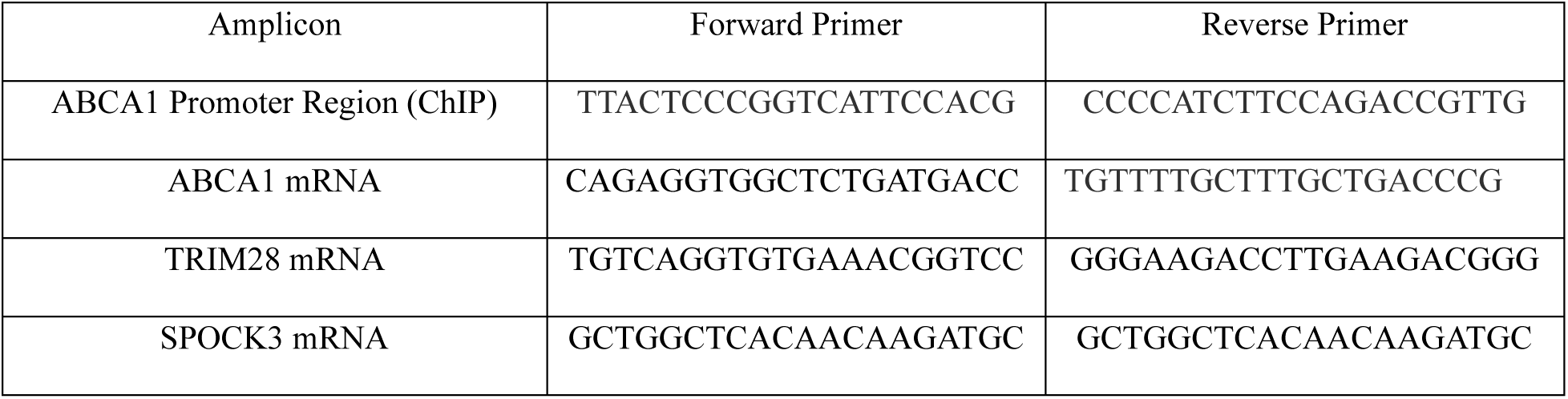
Primers used in the study.

### RNA Sequencing and Gene Expression Analysis

Single-end raw sequencing reads were first pre-processed by using fastqc (v0.10.1)[31] for sequencing base quality control. Reads were then mapped to RefSeq (RRID:SCR_003496) GRCh37-hg19 human reference genome and corresponding gene annotation obtained from UCSC’s repository using splicing-detection tools Bowtie (v1.0.1, RRID:SCR_005476)[32] and TopHat (v2.0.13, RRID:SCR_013035)[33] allowing a maximum of 1 mismatch per read. A second pass QC was done using alignment output with RSeQC (v2.6.3, RRID:SCR_005275)[34] to examine abundances of genomic features, splicing junction saturation and gene-body coverage. Gene expression is quantified using HTSeq[35] using the -m intersection-strict option. Differential expression analyses were performed using DESeq2 (v1.18.1)[36], a variance-analysis package developed to infer and detect differential gene expression in RNA-seq data. Downstream and visualization plots were done using regularized-log2 transformation implemented by DESeq2.

### Chromatin Immunoprecipitation (ChIP) qRT-PCR

Cells were grown in 10 cm dishes to 90-95% confluence. Two dishes were used per ChIP. Fixation was performed in the dish. 9.5 mL of PBS with 1% formaldehyde was added to each plate and incubated for 10 minutes with rocking. Fixation was stopped by adding 0.5 mL of 2.5M glycine for 5 minutes. Plates were washed twice with ice-cold PBS then 1 mL of PSB with protease inhibitors (Roche) was added to each dish. Cells were mechanically harvested using cell scrapers into 15 mL tubes, spun down into cell pellets, and lysed for 10 minutes in 300 μL Szak’s RIPA buffer (150 mM NaCl, 1.0% NP-40, 0.1% SDS, 50 mM Tris-HCl pH 8, 5 mM EDTA, 0.5 mM PMSF) with protease inhibitors. Following lysis, chromatin was sheared in a Bioruptor sonicator (Diagenode) in 1.5 mL TPX microtubes for Bioruptor (Diagenode, Cat No C30010010) for 45 seconds on/45 seconds off x 16 cycles on the high setting. Samples were spun for 10 minutes to clear debris, combined per condition, mixed well, and aliquoted for ChIPs. Input samples were set aside. ChIP samples were brought up to 1 mL with Szak’s RIPA buffer, and the appropriate amount of antibody was added per ChIP (STAG2: 3.5 μg (Cell Signaling Cat No. 5882), TRIM28: 2 μg (Proteintech Cat No. 15202-1-AP), Rabbit IgG: 1 μg (Invitrogen, Cat No. 1534782), H3K9ac: 10 μL (Cell Signaling Cat No. 9649S), H3K27ac: 5 μL (Cell Signaling, 8173S)). Samples were incubated overnight at 4C with rotation. The next day, ChIP-grade magnetic beads (ThermoFisher Scientific Cat # 88802) were washed 2x with Szak’s RIPA and 20 μL washed beads were added to each ChIP. Samples were incubated for 2 hours at 4C with rotation, followed by washes with low salt wash buffer (1% SDS, 1% Triton-X, 200 μM EDTA, 20 mM Tris-HCl pH 8, 500 mM NaCl), high salt wash buffer (1% SDS, 1% Triton-X, 200 μM EDTA, 20 mM Tris-HCl pH 8, 150 mM NaCl), LiCl wash buffer(0.0105975% LiCl, 1% NP-40, 1% sodium deoxycholate, 100 μM EDTA, 10 mM Tris-HCl pH), and two washes with TE buffer (100 μM EDTA, 10 mM Tris-HCl pH 8).

250 μL of elution buffer (70 mM Tris-HCl pH 8, 1.0 mM EDTA, 1.5% SDS) was then added to each sample, and inputs were brought up to 250 μL with elution buffer. Samples were eluted by incubating at 65C for 10 minutes then 95C for one minute. Supernatants were transferred to new tubes and reverse crosslinked by adding 13 μL of 4M NaCl and incubating overnight at 65C. The next day, 20 μg Proteinase K (Life Technologies Cat #EO0491) was added to each sample and incubated at 45C. DNA cleanup was performed utilizing the Monarch PCR & DNA Cleanup Kit (Cat # T1030S), and the final DNA was eluted in 50 μL water. 0.5-1 μL of ChIP DNA was used per qRT-PCR reaction.

### ChIP-sequencing

Samples were prepared for sequencing using the MAGnify™ Chromatin Immunoprecipitation System (Cat # 4392024, Thermo Fisher Scientific) for STAG2 with species-matched negative control IgG. 2 ng chromatin-immunoprecipitated DNA was used to generate the library for next-generation sequencing using the ThruPLEX DNA seq kit (Rubicon Genomics, Inc.) according to the manufacturer’s instructions. The DNA libraries were quantitated using the KAPA Biosystems qPCR kit and pooled in an equimolar fashion. Each pool was denatured and diluted to 2.4 pM with 1% PhiX control library. The resulting pool was loaded into the appropriate NextSeq Reagent cartridge for 75 paired-end sequencing and sequenced on a NextSeq500 following the manufacturer’s recommended protocol (Illumina). ChIP-Seq reads were mapped to reference genomes using bwa [37], and the narrow peaks were identified by MACS2 [38] using the input DNA as a control.

### Rapid Immunoprecipitation with Mass Spectrometry of Endogenous Proteins

#### Antibody conjugation to beads

Magnetic beads were vortexed and aliquoted (100 µl) into 2 ml round-bottom tubes placed on a chilled magnetic stand. Beads were washed and blocked four times with 1 ml PBS containing 5 mg/ml BSA (PBS/BSA). After resuspension in 500 µl PBS/BSA, the appropriate antibody (STAG2 and species-matched IgG) was added, and the mixture was rotated either overnight at 4 °C or for 1 h at room temperature. Antibody-bound beads were subsequently washed five times with 1 ml PBS/BSA to remove unbound antibody.

#### Cell crosslinking and cell lysis

T24 cells (2 × 10^7) were crosslinked in 10 ml serum-free medium containing 1% formaldehyde for 8 min at room temperature. The reaction was quenched with 1 ml of 1 M glycine (final concentration 0.1 M), and cells were washed twice with 10 ml ice-cold PBS. Cells were collected with a silicone scraper in 500 µl PBS per plate, pelleted at 2,000 × g for 3 min at 4 °C, and washed once more in 500 µl PBS. Cell pellets were sequentially lysed in 10 ml LB1 (50 mM HEPES-KOH, (pH 7.5), 140 mM NaCl, 1 mM EDTA, 10% (vol/vol) glycerol, 0.5% (vol/vol) NP-40/Igepal CA-630 and 0.25% (vol/vol) Triton X-100) (10 min rotation at 4 °C) and 10 ml LB2 (10 mM Tris-HCL (pH 8.0), 200 mM NaCl, 1 mM EDTA and 0.5 mM EGTA)(5 min rotation at 4 °C), with centrifugation at 2,000 × g for 5 min between steps. Nuclear pellets were resuspended in 300 µl LB3 (10 mM Tris-HCl (pH 8.0), 100 mM NaCl, 1 mM EDTA, 0.5 mM EGTA, 0.1% (wt/vol) sodium deoxycholate and 0.5% (vol/vol) *N*-lauroylsarcosine) and sonicated on ice (30 s on/off cycles for 10 min) to yield DNA fragments of 200–600 bp. Lysates were clarified by addition of 30 µl 10% Triton X-100 and centrifugation at 20,000 × g for 10 min at 4 °C.

#### Immunoprecipitation

One hundred microliters of antibody-conjugated beads were added to clarified lysates and rotated overnight at 4 °C. Beads were washed ten times with 1 ml RIPA buffer, followed by two washes with 1 ml freshly prepared 100 mM ammonium bicarbonate. For the second ammonium bicarbonate wash, beads were transferred to new tubes.

#### On-bead digestion and peptide desalting

Bead-bound proteins were digested directly with 10 µl (10 ng/μL) of sequencing-grade trypsin diluted in 100 mM ammonium bicarbonate. Samples were vortexed intermittently during the first 15 min and incubated overnight at 37 °C. A second aliquot of 10 µl trypsin (10 ng/µl) was added the following day for an additional 4 h digestion. Digested peptides were recovered from the supernatant (∼20 µl) and acidified to a final concentration of 5% formic acid by adding 1 µl 100% formic acid.

Peptides were desalted using C18 ultra-micro spin columns. Cartridges were conditioned with 2 × 100 µl 50% acetonitrile/water and equilibrated with 2 × 100 µl 0.1% formic acid. Acidified peptides were loaded, reloaded up to three times to maximize binding, washed four times with 0.1% formic acid, and eluted twice with 50 µl 60% (v/v) acetonitrile/0.1% (v/v) formic acid. Eluates (∼100 µl total) were pooled, dried by vacuum centrifugation, and stored at –20 °C or – 80 °C until LC-MS/MS analysis.

#### LC-MS/MS acquisition

Peptides were reconstituted in 0.1% formic acid. The digests were analyzed by nanoLC-MS/MS in a ThermoScientific Q-Exactive Mass Spectrometer coupled to a nanoHPLC EASY-nLC 1000 (ThermoScientific). For the LC ESI–MS/MS analysis, ∼1 μg of peptides was loaded onto the column and eluted for 120 min using a reverse-phase column (C18, 2 μm, 100 A, 50 μm × 150 mm), Easy-Spray Column PepMap RSLC (P/N ES801)) suitable for separating protein complexes with a high degree of resolution. The flow rate used for the nano column was 300 nl min-1 and the solvent gradient range was 7% B (for 5 min) to 35% B in 120 min. Solvent A was 0.1% formic acid in water whereas B was 0.1% formic acid in acetonitrile. The injection volume was 2 μl. The MS equipment has a high collision dissociation cell (HCD) for fragmentation and an Orbitrap analyzer (Q-Exactive-ThermoScientific Germany). A voltage of 3.5 kV was used for Electro Spray Ionization (ThermoScientific, EASY-SPRAY). XCalibur 3.0.63 (ThermoScientific) software was used for data acquisition with a configuration that allows peptide identification at the same time as their chromatographic separation. A Data dependent method was used: Full-scan mass spectra were acquired in the Orbitrap analyzer. The scanned mass range was 400–1800 m/z, at a resolution of 70000 at 400 m/z and the twelve most intense ions in each cycle were sequentially isolated, fragmented by HCD, and measured in the Orbitrap analyzer. Peptides with a charge of +1 or with an unassigned charge state were excluded from fragmentation for MS2.

#### Analysis of LC ESI–MS/MS data

Raw data generated with Xcalibur software was processed and analyzed with Proteome discoverer 2.1.1.21 with SEQUEST Search engine. Spectrum Selector node with default parameter settings was used to generate peak lists. Minimum and maximum precursor masses were set at 350 and 5000 with an S/N of 1.5. Data were searched against Uniprot Homo sapiens database, with trypsin specificity (full cleavage) and a maximum of two missed cleavages per peptide. Carbamidomethylation of cysteine residues was set as a fixed modification and oxidation of methionine was set as variable modification. Proteome Discoverer searches were performed with a precursor mass tolerance of 10 ppm and product ion tolerance to 0.05 Da. Proteome Discoverer default settings were used: Target FDR = 0.01; Z = 1 High confidence XCorr 1.5; Z = 2 High confidence XCorr 2; Z = 3 High confidence XCorr 2.5; z ≥ 4 High confidence XCorr 3. Protein hits were filtered for high confidence peptide matches with a maximum protein and peptide false discovery rate of 1% calculated by employing a reverse database strategy.

### Cistrome-BETA Functional Predication Analyses

DEGs from RNAseq DESeq2 analyses and .bed files from STAG2 ChIP-seq were uploaded to the Galaxy / Cistrome server for functional prediction analysis (http://cistrome.org/ap/root) [39].

BETA-plus analysis was performed with the following parameters: Peaks considered to contribute to the genes: 10,000; the distance from gene TSS within which peaks will be selected: 100,000; padj < 0.05, degree of gene expression change (log2FC) > 0.1375.

### Hi-C Sample Preparation and Sequencing

T24 cells were cultured as previously described, harvested by trypsinization, and counted. 1*10^6^ cells were aliquoted and pelleted. 1*10^6^ cells were used per Hi-C reaction. Sample preparation was performed using the Arima Genomics Hi-C+ kit (Cat No. A510008). All steps were carried out according to the manufacturer’s protocol for standard input samples, including all quality control steps (Arima-Hic+ Kit A160509 User Guide). Library preparation was performed by the Roswell Park Genomics Shared Resource using the Arima Library Prep Kit (Cat No. A303011) according to manufacturer’s protocol. Final purified libraries are validated for appropriate size on a 4200 TapeStation D1000 Screentape (Agilent Technologies, Inc.). The sequencing libraries are quantitated using KAPA Biosystems qPCR kit (Roche, Inc.) and are pooled together in an equimolar fashion, following experimental design criteria. The resulting pool is then loaded into the appropriate NovaSeq Reagent cartridge and sequenced on a NovaSeq6000 following the manufacturer’s recommended protocol (Illumina Inc.). 600 paired-end reads were sequenced for each condition in duplicate. Duplicates within conditions were pooled for downstream analyses.

### Hi-C Computational Analysis

#### Preprocessing: Alignment, Filtering, Binning

The raw sequencing reads were preprocessed using runHiC (v0.9.0). Briefly, paired-end reads were aligned to the human reference genome (hg38) using chromap(v0.3.1) as the aligner. Mapping was performed using gzip-compressed Hi-C FASTQ files, with read identifiers retained and dangling-end and other noninformative reads removed. Alignment was carried out using a chunk size of 1,500,000 bp, with a total of 10 processing threads and 64 GB of memory allocated. Following alignment, filtering, and quality control procedures, Hi-C contacts were binned using runHiC with default parameters. The binning step was performed using 10 parallel processes, producing filtered and binned interaction files at resolutions of 5 kb, 10 kb, 25 kb, 50 kb, 100 kb, 250 kb, 500 kb, 1 Mb, and 2.5 Mb for downstream analysis.

#### Contact Frequency Matrix Generation

Downstream Hi-C analyses were conducted using cooltools (v0.7.1). Genome-wide Hi-C contact frequency matrices, representing the intensity of interaction between different genomic regions in the same chromosome, were generated for cis-interactions and analyzed at a resolution of 100kb.

#### Compartment Analysis

A/B compartments were identified by eigenvector decomposition of the observed contact matrices. The first eigenvector (E1) was used to assign genomic bins to compartments, with positive and negative values corresponding to A and B compartments, respectively. Compartmentalization strength was assessed using saddle plots, restricting the eigenvector quantile range to 0.02–0.98.

#### Insulation/Boundary Analysis

Topologically Associating Domains (TADs) are the basic units of the chromatin structure and are identified by local enhancement regions. The TAD organization was evaluated by calculating insulation scores with sliding window sizes of 30 kb and 50 kb at 10-kb resolution, restricted to valid genomic regions defined by an hg38 view file. Domain boundaries were identified by applying Li’s minimum cross-entropy thresholding method to the insulation score profiles.

#### Contact Length Quantifications

Contact length changes were assessed between STAG2 KO and control conditions by calculating the distance between the start of region 1 and the start of region 2 for paired regions with significant (adjusted p-value < 0.05) changes in contact frequency (absolute log2FC difference >1). Comparisons of genomic distance between pairs were made between pairs which gained contact frequency (positive log2FC) and pairs which lost contact frequency (negative log2FC).

#### Chromatin Loop Detection and Focal Enrichment

Chromatin interaction loops (dots) were identified from balanced Hi-C contact matrices using the cooltools package. Hi-C contact maps were stored in multi-resolution cooler format and analyzed independently for each condition (A6, G2, and H2) at 25-kb resolution using the human genome (hg38). Chromosome sizes and centromere annotations were obtained from the USCS Genome Browser through bioframe library, and chromosome arms were generated to define genomic regions.

To account for the distance-dependent decay of chromatin contacts, cis-expected contact frequencies were calculated for each chromosome arm. Expected values were estimated from balanced contact matrix and used as local background models for loop detection. Significant focal interaction enrichments (dots) were subsequently identified using the cooltools.dots function, which compares observed contact frequencies with locally adjusted expected interaction frequencies derived from the multiple neighborhood kernels (donut, horizontal, vertical, and lower-left), following an approach conceptually similar to HiCCUPS. Dot calling was restricted to intra-chromosomal interactions separated by no more than 10 Mb.

To visualize focal interaction enrichment, balanced observed/expected (O/E) contact matrices were generated and displayed as heatmaps, and focal enrichment patterns were examined across the A6, G2, and H2 conditions using identical visualization parameters. The resulting dot calls were used to characterize condition-specific chromatin looping and local interaction enrichment landscapes.

### Regulatory Transcriptional Network (RTN) Analysis

STAG2-specific differentially expressed genes (DEGs) from the Roswell Park patient cohort were utilized for regulatory transcriptional network (RTN) analysis utilizing the Bioconductor RTN package [40, 41]. Normalized expression data generated from DeSeq2 [36] were used as expression data, and the identified STAG2-specific DEGs were analyzed for TRIM28 regulatory potential. 2-tailed gene set enrichment analysis was performed with 100 permutations to detect enrichment for TRIM28 regulatory potential in up- or down-regulated DEGs. Plots were generated using the tna.plot.gsea2 function within the RTN package.

### ShinyGO Pathway Analysis

Bioconductor package GenomicRanges version 1.58.0 was utilized to annotate regions of the genome experiencing compartment shifts (A to B, or B to A) with genes contained within each region. In either compartment shift category (A to B, or B to A), only genes identified from RNA-seq of T24 STAG2 KO vs. Control cells were selected for further analysis. Gene lists for A to B regions and B to A regions were generated and input into the ShinyGO 0.85 web interface (http://bioinformatics.sdstate.edu/go/) [42] and analysis was performed for GO:Biological Pathways with default settings. Pathways were considered significantly enriched in either gene set if the Enrichment FDR was < 0.05. Pathways were plotted using ggplot in R and ranked based on Fold Enrichment.

### Additional Statistical Analyses

Quantitative measures are compared between two groups using two-sample t-tests, three groups via one-way ANOVA, and data with multiple groupings via two-way ANOVA in GraphPad Prism unless otherwise noted. A *P* value of 0.05 was set to be statistically significant. Statistical analyses for genomics experiments were performed as described in previous sections.

## RESULTS

### STAG2 transcriptionally regulates pro-tumor pathways in bladder cancer and promotes invasion

To define the transcriptional landscape of bladder cancer, we performed RNA-sequencing of tumor (T; n = 66) and adjacent non-tumor (N; n = 15) bladder samples obtained from Roswell Park Comprehensive Cancer Center (RP cohort). Differential expression analysis followed by Ingenuity Pathway Analysis (IPA) revealed significant enrichment of the cohesin chromatin regulation pathway in tumor samples compared to non-tumor controls, highlighting a potential role for cohesin-mediated chromatin regulation in bladder cancer tumorigenesis and progression (Figure S1A, Table S1). The cohesin member STAG2 acts in an oncogenic manner in bladder cancer, as previously shown by our group and others [10, 11], and therefore we sought to identify STAG2-specific tumor characteristics in the bladder cancer setting. To characterize transcriptional profiles associated with STAG2 protein expression we compared gene expression profiles between groups defined by STAG2 tumor protein expression (Figure S1B), based on previously established cutoff values [10]. We found 83 differentially expressed genes (DEGs) between the STAG2-low (n= 38) and STAG2-high (n = 17) cohorts (Figure 1A, Table S2). Performing pathway analysis of Hallmark gene signatures from MSigDB [43] revealed that STAG2 high tumors were transcriptionally enriched for the epithelial to mesenchymal transition (EMT) pathway in addition to enrichment of several other pro-tumor pathways such as MYC targets and TGFβ signaling (Figure 1A, right).

**Figure 1.**
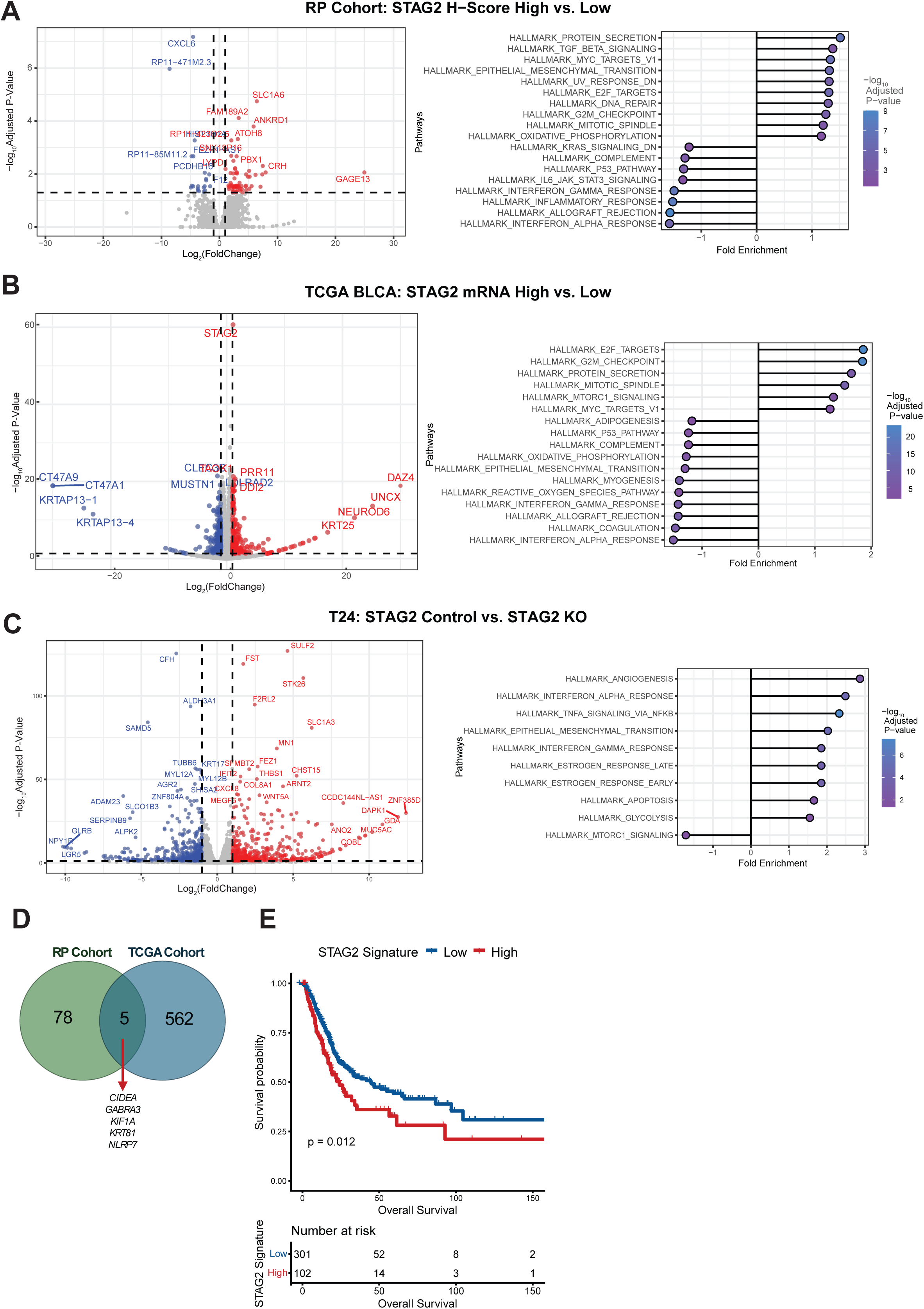
STAG2 transcriptionally regulates pro-tumor pathways in bladder cancer and promotes invasion. A. Left: Differentially expressed genes (DEGs) between patient tumors with high STAG2 expression (H-score > 50) and patients with low STAG2 expression (H score < 50). Positive log2 Fold Change (Log2FC) indicates gene is downregulated in STAG2-low tumors, negative log2FC indicates gene is upregulated in STAG2-low tumors. Right: Pathway analysis utilizing Hallmark gene sets from MSigDB based on the identified DEGs. B. Left: DEGs from TCGA BLCA STAG2-high (n=204) and STAG2-low (n=204) cohorts, stratified based on median STAG2 expression. Positive log2FC indicates gene is upregulated in STAG2-high tumors, negative log2FC indicates gene is upregulated in STAG2-low tumors. Right: Pathway analysis utilizing Hallmark gene sets from MSigDB based on the identified DEGs. C. Left: Volcano plot indicating DEGs between T24 control (STAG2 WT) and STAG2 KO cells. Positive log2FC indicates gene is upregulated in the control setting, negative log2FC indicates gene is upregulated after STAG2 KO. Right: Pathway analysis utilizing Hallmark gene sets from MSigDB based on the identified DEGs. D. Venn diagram of DEGs from RP and TCGA cohorts determined by median STAG2 mRNA expression, with five genes overlapping between both datasets. E. Kaplan-Meier survival curve for the TCGA patient cohort based on STAG2 signature score, comparisons made between the top 25% signature scored patients (n = 102) versus the bottom 75% (n = 301) signature scored patients. P-value determined by log rank test.

To extend these findings to a larger cohort of MIBC patients, we performed differential expression analysis based on STAG2 mRNA expression in the TCGA BLCA cohort [29] (Table S3). Notably, STAG2 mRNA expression showed a modest but significant correlation with STAG2 protein levels (H-score) in the RP cohort (Figure S1C), supporting the use of STAG2 mRNA expression as an adequate surrogate marker for STAG2 protein abundance in downstream analyses. Thus, we divided patients into *STAG2*-high and *STAG2*-low cohorts based on median expression of *STAG2* mRNA. We identified 567 DEGs based on *STAG2* mRNA levels, with 262 upregulated in the *STAG2*-high cohort and 305 upregulated in the *STAG2*-low cohort (Figure 1B). Pathway analysis revealed differential enrichment of similar pathways as identified in the RP cohort, such as MYC targets, and several immune-related pathways (Figure 1B).

To determine whether STAG2-specific gene expression programs identified in clinical samples would be reflected *in vitro,* we performed RNA sequencing (RNA-seq) of one clonally generated T24 MIBC cell line with wild-type (WT) expression of STAG2 and two CRISPR-induced T24 STAG2 knock out (KO) cell lines. These cell lines recapitulate our previous findings in which STAG2 expression is significantly associated with an invasive phenotype (Figure S2A) [10]. 481 genes were upregulated and 421 genes were downregulated in the STAG2 WT setting (Figure 1C, Table S4). Many cancer-related pathways were transcriptionally enriched in the STAG2 WT setting, including the epithelial to mesenchymal transition (EMT) pathway, reflecting clinical sample analysis, in addition to several immune-signaling related pathways such as interferon alpha response, TNFα signaling via NFKβ, and interferon gamma response (Figure 1C).

MIBC patients with high STAG2 protein expression experience worse survival outcomes [10], however when stratifying patients by *STAG2* mRNA expression level, we do not see a difference in survival (Figure S1D). We speculate that the activity of STAG2, rather than just mRNA expression level, may influence clinical outcomes. To investigate the contribution of STAG2 activity level to patient survival outcomes, we generated a STAG2 activity signature based on common DEGs identified between the RP cohort and the TCGA cohort after stratifying by *STAG2* mRNA expression level (Figure 1D) and utilized these genes to generate a STAG2 activity score for each individual patient. Survival analyses of patients in the TCGA cohort demonstrate that patients with the top quartile of STAG2 activity score suffer significantly shorter overall survival compared to remaining 75% of patients (Figure 1E). We did not see a difference in overall survival between STAG2 activity-high and STAG2 activity-low cohorts within the RP dataset, which may be due to the small sample size (Figure S1E). Altogether, these data emphasize the importance of investigating mechanisms of STAG2-mediated gene regulation in MIBC to dissect how STAG2 contributes to pro-tumor transcriptional programs, as these may have consequences on patient survival.

### STAG2 directly and indirectly restricts aberrant chromatin contacts driving a gene regulatory function in MIBC cells

The cohesin complex is essential for maintenance of chromatin organization and subsequent regulation of gene expression. Consequently, we sought to identify how STAG2-mediated chromatin regulation may contribute to the pro-oncogenic phenotype associated with STAG2 in bladder cancer. To assess chromatin organization and characteristics, we performed genome-wide Hi-C in the previously generated T24 control and STAG2 KO cell lines. We generated contact frequency matrices for each condition, denoting regions of high and low probability of chromatin contacts which can be visualized at the whole-genome level (Figure S2B), individual chromosome level (Figure S2C), or single-gene level.

To comprehensively analyze contact frequency alterations after STAG2 KO, we quantified the number of regions with statistically significant gains or losses in contact genome-wide (Figure 2A, S2D, Table S5). When separated by chromosome, we found that chromosome 4 experienced the largest degree of contact gain, and chromosome 9 experienced the largest degree of contact loss (Figure S2E). An example denoting changes in chromatin contact for a 3Mb region of chromosome 9, which contains several protein-coding genes, is shown in Figure 2B. We next asked whether the changes in contact of chromatin regions were longer-ranged or shorter-ranged based on whether contact was gained or lost. The distance between regions gaining contact with each other was significantly longer than the distance between regions which lost contact, indicating a gain of long-range contacts and a collapse of short-range contacts after STAG2 KO (Figure 2C). This is consistent with reports suggesting that STAG2-containing cohesins maintain shorter-ranged contacts, whereas STAG1-containing cohesins maintain longer-ranged contacts [44]. To evaluate the impact of altered chromatin contact on gene expression, we identified DEGs from RNA-seq which are encoded in regions that gained or lost contact with other regions in the genome. We found a significant association between the magnitude of gene expression change and the direction of contact change, where 88% of DEGs experiencing a gain-of-contact were upregulated after STAG2 KO, and 55% of DEGs experiencing a loss-of-contact were downregulated (Figure 2D, p = 0.0011, Fisher’s exact test). To ask whether changes in contact were direct effects of STAG2 loss, we analyzed the proportion of STAG2-bound regions that had altered contact frequency. After integrating ChIP-seq data for STAG2 binding, we identified only a small proportion of sites (1.77%) that were STAG2 bound at baseline, suggesting that contact changes post-STAG2-KO are not simply due to loss of STAG2 occupancy on DNA (Figure 2E). However, the distance between STAG2-bound regions that gained contact frequency was significantly longer than STAG2-bound regions that lost contact frequency (Figure 2F). This suggests that STAG2 directly mediates short-ranged genomic contacts.

**Figure 2.**
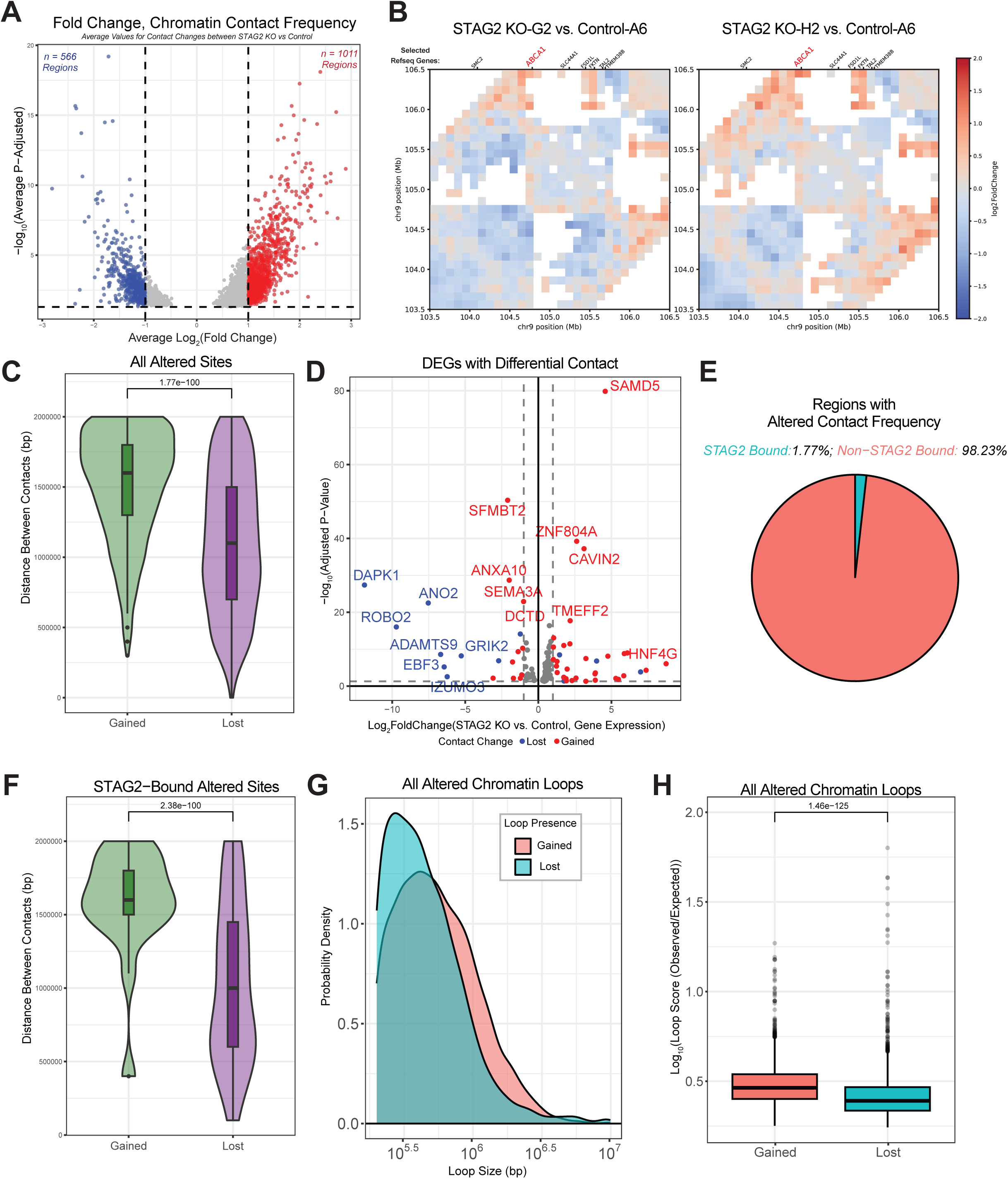
STAG2 directly and indirectly restricts aberrant chromatin contacts driving a gene regulatory function in MIBC cells. A. Volcano plot indicating the log2FC for contact between pairs of genomic regions of STAG2 KO versus control cells. One point represents one pair of genomic regions. Positive log2FC indicates contact between regions is increased after STAG2 KO, negative log2FC indicates contact between regions is decreased after STAG2 KO. B. Heatmap showing regions of statistically significant contact frequency changes (indicated by log2FC) for an example range on chromosome 9. RefSeq genes encoded in this region are shown at the top. Left, STAG2 KO-G2 versus control, right, STAG2 KO-H2 versus control. Blue indicates loss of contact, red indicates gain of contact after STAG2 KO. C. Distance between all genomic pairs that gained contact after STAG2 KO (green) or lost contact after STAG2 KO (purple). P-value determined by Wilcoxon signed-rank test. D. Dots represent DEGs encoded in regions of differential contact frequency; red experienced increase in contact, blue experienced loss of contact. E. Pie chart of all regions that experienced altered contact, separated into STAG2 bound and non-STAG2 bound regions. F. Distance (base pairs) between genomic pairs containing one or more STAG2 binding sites that gained contact after STAG2 KO (green) or lost contact after STAG2 KO (purple). P-value determined by Wilcoxon signed-rank test. G. Density plot of loop size (measured in base pairs) for chromatin loops gained after STAG2 KO or lost after STAG2 KO. H. Log10-transformed loop score for loops gained and lost after STAG2 KO as assessed by the observed over expected value of the enriched contact point from Hi-C contact maps. P-value determined by Wilcoxon signed-rank test.

Individual chromatin loops can be inferred from Hi-C data by identifying “dots,” or punctate foci with enriched chromatin contact frequency on contact frequency heatmaps. To identify differential chromatin loops between the control and STAG2 KO settings, we identified, quantified, and compared the number and genomic location of enriched foci in STAG2 KO cells versus control (Table S6). Consistent with changes in global chromatin contacts, we found that loops lost after STAG2 KO were smaller, and that loops gained in the STAG2 KO setting were longer (Figure 2G). Based on the observed over expected value of each enriched foci, we next inferred the strength of the chromatin associations forming loops. Chromatin loops that emerged after STAG2 KO had a greater observed over expected loop score than chromatin loops that were lost after STAG2 KO, suggesting that loss of STAG2 leads to longer ranged, more stable chromatin looping genome-wide (Figure 2H).

We can infer changes in regions of genomic insulation, or “boundaries” separating the genome into different domains, from Hi-C data. Given the role of STAG2-cohesin in maintaining short-range genomic contacts [44], we hypothesized that STAG2 KO would cause loss of insulation. To characterize insulator regions in each condition we calculated insulation scores genome-wide at 10kb resolution, and to identify regions of altered insulation between STAG2 KO and control cells. Overall, STAG2 KO resulted in a net loss of insulation genome-wide (8,253 regions in control, 8,150 and 8,096 in KO G2 and H2, respectively; Table S7). When segregated by chromosome, however, there were chromosome-specific effects. Some chromosomes experienced a net gain of insulator regions after STAG2 KO, other chromosomes experienced a net loss of insulator regions after STAG2 KO. Chromosome 7 had the largest magnitude of insulation gain (429 regions in control, 441 and 439 in KO-G2 and -H2, respectively), whereas chromosome 9 had the largest magnitude of insulation loss (343 regions in control, 327 and 318 in KO-G2 and -H2, respectively).

Together, our Hi-C results support the hypothesis that STAG2 directly and indirectly supports chromatin organization, loop formation, and domain formation with observable consequences on gene expression that may underlie the pro-oncogenic function of STAG2 in MIBC.

### Chromatin compartment shifts drive changes in gene expression patterns after STAG2 KO

Chromatin compartmentalization is an aspect of chromatin organization that can have significant consequences on magnitude of gene expression [22, 23]. Chromatin in the A compartment tends to be more transcriptionally active, more accessible, and localized to the interior of the nucleus [23]. Conversely, chromatin within the B compartment is more less transcriptionally active and localized to the nuclear periphery [23]. Therefore, we next assessed whether changes in chromatin organization post-STAG2-KO result in altered chromatin compartmentalization. To quantify chromatin compartment changes across the whole genome, we segmented the linear genome into 100 kb bins and evaluated the compartment score of each 100 kb region (as assessed by the first principal component). Then, we compared compartment score of the control cell line regions with both STAG2 KO cell lines. Overall, most regions maintained their compartmentalization after STAG2 KO (78.3% for STAG2 KO-G2, 81.5% for STAG2 KO-H2 vs control-A6; Figure 3A, Table S8). However, a small percentage of regions went from A in control to B in the STAG2 KO setting (2.7% for A6-G2 comparison and 3.3% for A6-H2 comparison) or conversely went from B in the control setting to A in the STAG2 KO setting (9.2% for A6-G2 and 5.7% for A6-H2; Figure 3A). Compartment states were relatively consistent across chromosomes (Figure S3A). Interestingly, the magnitude of B to A transitions was larger than A to B in both comparisons. This supports a region-specific function of STAG2 in maintaining both A and B compartment chromatin, with a slightly more dominant role in maintaining less active chromatin.

**Figure 3.**
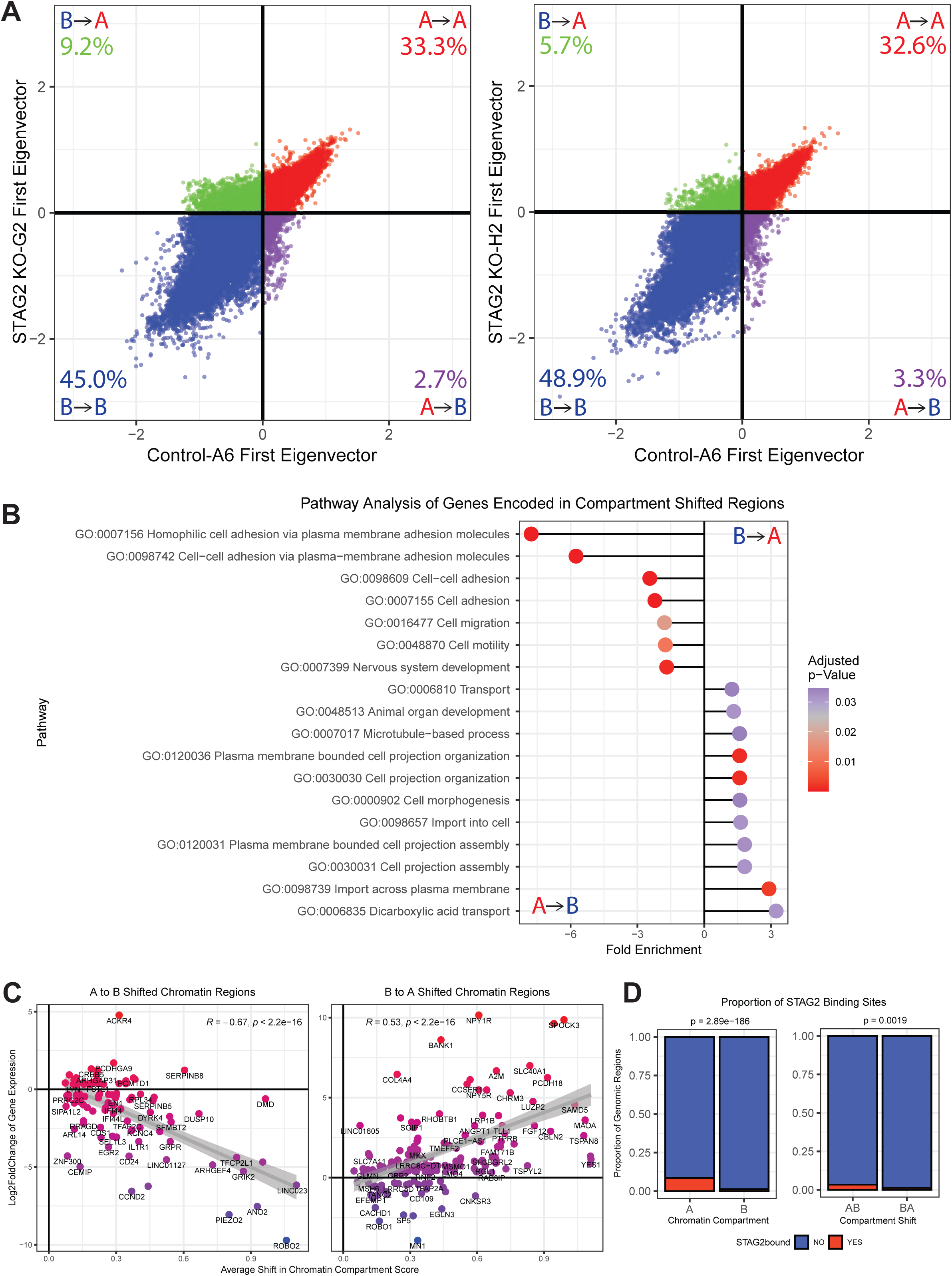
Chromatin compartment shifts drive changes in gene expression, increase enrichment of apoptotic pathways, and decrease enrichment of motility pathways after STAG2 KO. A. Scatter plots depicting the first principal component score for each 10kb region across the genome, with positive score indicating compartment A, and negative score indicating compartment B for comparisons of T24 control A6 (STAG2 WT) versus T24 STAG2 KO-G2 (left), and T24 control A6 versus T24 STAG2 KO-H2 (right). Percentage of 10kb-bins in each category (A in control and KO, B in control and KO, A in control and B in KO, and B in control A in KO) displayed. B. ShinyGO Pathway Analysis of GO: Biological Process pathways for genes encoded in regions shifting from compartment A to B (negative fold enrichment) and regions shifting from compartment B to A (positive fold enrichment). C. Scatter plot of average shift in compartment score and log2FC for all DEGs experiencing compartment shifts. Each dot is representative of a single gene. Correlation plotted using the geom_smooth function in R, Pearson correlation coefficient and p-value displayed. D. Proportion of A and B compartments (left) or A to B and B to A compartment shifts (right) colored by proportion of STAG2 binding sites. P-values assessed using Fisher’s exact test.

To determine whether changes in chromatin compartment score corresponded with changes in gene expression, we identified the protein-coding genes contained in each 100 kb bin that had been assigned a compartment score within the genome. We generated two gene lists: one containing all genes encoded in regions shifting from A to B after STAG2 KO, and one containing all genes encoded in regions shifting from B to A. Utilizing ShinyGO [42], we performed pathway enrichment analysis on both gene lists to identify enriched biological processes in either setting. Genes contained in regions shifting from A to B compartments were significantly enriched for cell migration and motility biological processes (Figure 3B). As B-compartment chromatin is less transcriptionally active, A to B shifts may result in downregulation of migration and motility gene sets, explaining the diminished invasive potential that we observe phenotypically after STAG2 KO (Figure S2A). Genes contained in regions shifting from B to A compartments were cell projection processes, as well as molecule transport pathways (Figure 3B). This demonstrates a broad role for chromatin compartmentalization in regulating gene expression programs related to cell migration and invasion. To assess whether the degree of compartment shift impacts the magnitude of observed gene expression changes, we performed correlation analysis of average shift in compartment score and average change in gene expression score for every shifted region containing a DEG after STAG2 KO. In A to B shifted regions, there was a significant negative correlation identified, and for B to A shifted regions, there was a significant positive correlation identified, implicating a strong impact and directional impact of compartment state on gene expression level (Figure 3C).

Finally, we asked whether these effects were directly due to loss of STAG2 binding at specific regions in the genome. To evaluate STAG2 localization, we performed ChIP-sequencing for STAG2 in T24 cells and integrated the location of the identified STAG2 binding peaks with the compartment score for each region. When looking at individual chromosomes, regions bound by STAG2 were largely A compartment-like and remained so after STAG2 KO (Figure S3B). In line with this, we found a statistically significant association of STAG2 binding in the A compartment at baseline, and that STAG2 binding is more frequent in regions shifting from A to B rather than B to A (Figure 3D). However, when broken down by chromosome, STAG2-bound regions experienced compartment shifts in both directions in a chromosome-dependent manner (Figure S3C). From our global compartment analysis, we identified more regions shifting from B to A (Figure 3A). Given the propensity of STAG2 binding in A and A to B regions, it is likely that the shift of a larger majority of B to A chromatin regions is not simply due to loss of STAG2 binding, but an effect of larger-scale genome reorganization that occurs after STAG2 loss. To evaluate the impact on gene expression of STAG2 bound genes in compartment shifted regions, we assessed correlation between compartment shift and gene expression and found a negative correlation between STAG2-bound gene expression and magnitude of compartment shift (Figure S3D). Altogether, this data indicates that STAG2 has direct and indirect roles in regulating chromatin compartmentalization which contribute to the overall role of STAG2 in mediating gene expression.

### STAG2-mediated compartmentalization restricts regulatory elements to modulate gene expression

Cohesin complexes can act as a bridge between genomic regions to maintain proximity of regulatory elements in physical space [45]. To determine whether altered contact of genes with regulatory elements is driving gene expression changes in T24 cells post-STAG2 KO, we integrated our Hi-C data with candidate cis-regulatory elements (cCREs) available from ENCODE [46], including promoter like (PLS), proximal-enhancer-like (pELS), distal-enhancer-like (dELS), DNAse-H3K4me3 (active promoter), and CTCF regulatory elements. We isolated STAG2-specific DEGs from our *in vitro* RNA-seq analysis, then evaluated which genes had differential contact with CRE regions in the genome after STAG2 KO. In line with our global analyses showing a larger magnitude of contact gains compared to contact losses genome-wide, both upregulated and downregulated DEGs had increased contact with a variety of cCREs throughout the genome. The number of upregulated genes was larger in magnitude (n = 30 upregulated genes and n = 20 downregulated genes with increased cCRE contact; Figure 4A). Less DEGs lost contact with cCREs compared to gained contact with cCREs, regardless of directionality of expression; however, only four upregulated DEGs lost cCRE contact while nine downregulated DEGs lost cCRE contact (Figure 4A). The top seven downregulated genes all lost contact with regulatory elements, and nine out of the top ten upregulated genes gained contact with regulatory elements, indicating direct function of chromatin contact in promoting gene expression. The type of cCRE which DEGs gained or lost contact with varied widely between each DEG. The dominant cCRE with altered DEG contact frequency was distal enhancer (dELS, Figure 4A). When we stratified up- and downregulated genes, we found that upregulated genes gained contact with a larger proportion of promoter-like (PLS), proximal-enhancer like (pELS), and active promoter regions compared to downregulated genes (Figure 4B). Conversely, downregulated genes lost contact with a large proportion of regions containing both distal and proximal enhancers compared to upregulated genes (Figure 4B). This indicates that altered contact with regulatory elements due to loss of STAG2 may contribute to changes in gene expression. To evaluate whether STAG2-mediated chromatin compartmentalization may contribute to altered gene-cCRE contact, we evaluated the frequency compartment shifts by cCRE type. The proportion of B to A compartment shifts was larger than A to B after STAG2 KO across all cCREs, suggesting a global opening of chromatin at regulatory elements that may permit increased contact with genes (Figure S4A-B). To identify whether this is a direct effect of STAG2 binding, we evaluated STAG2 binding at each CRE region (Figure S4C) and found that STAG2 was most frequently found at PLS and dELS regions, in line with what has been reported previously (Figure S4C) [19]. Evaluation of compartment shifts at STAG2-bound CREs revealed that STAG2 maintains A compartment chromatin in most regions, except in the case of STAG2-bound dELS and CTCF-only sites, which were most likely to shift from compartment B to A (Figure 4C, S4 D-E).

**Figure 4.**
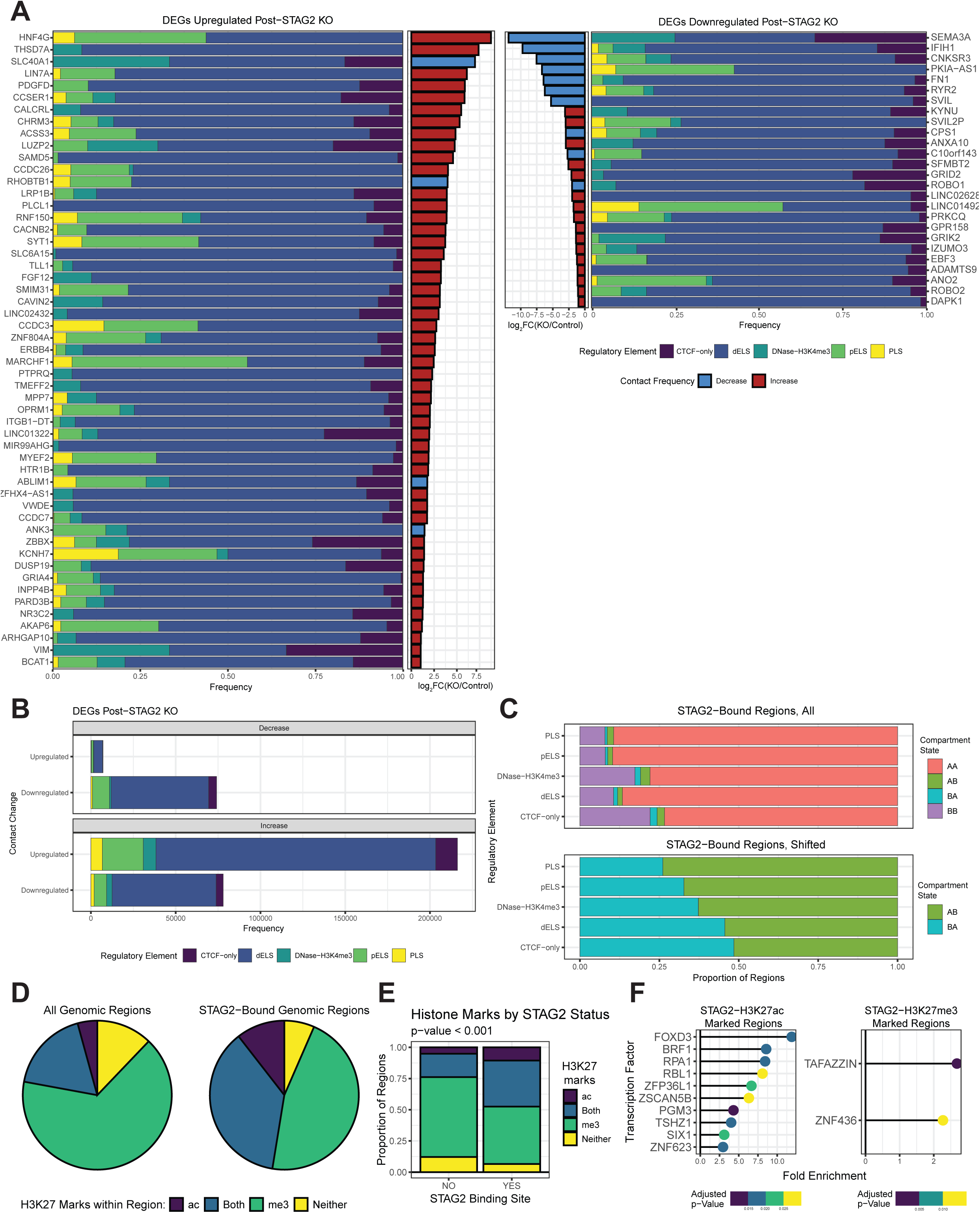
STAG2-mediated compartmentalization restricts regulatory elements to modulate gene expression. Frequency of altered contact with candidate cis-regulatory elements (cCREs) for upregulated DEGs (left) and downregulated DEGs (right), ordered by magnitude of change in gene expression (log2FC). CTCF-bound and non-CTCF-bound regions were combined for dELS, DNAse-H3K4me3, pELS, and PLS regions. B. Frequency of altered contact for DEGs with decreased cCRE contact (top) increased cCRE contact (bottom), with CTCF-bound and CTCF-unbound regions combined for dELS, DNAse-H3K4me3, pELS, and PLS regions C. Top: Proportion of compartment states for STAG2-bound cCREs: AA: A remaining A compartment; AB: A to B compartment switch; BA: B to A compartment switch; BB: B remaining B compartment. Bottom: Proportion of STAG2-bound, compartment shifted cCREs, only: A to B or B to A. D. Pie chart depicting distribution of regions containing H3K27ac, H3K27me3, both, or neither for the whole genome (left) or at only STAG2 bound regions (right). E. Proportion of STAG2-bound and non-STAG2-bound genomic regions containing H3K27ac, H3K27me3, both, or neither. P-value assessed using chi-squared test of independence. F. Enrichment of C3 MSigDB transcription factor target pathways for all genes encoded in regions denoted as STAG2 and H3K27ac-only (left) or STAG2 and H3K27me3-only (right).

Given the identified association of the cohesin complex with the PRC2 complex [25, 47] and the localization of STAG2 at promoters and enhancers, we next asked whether STAG2 is more likely to localize to regions containing H3K27ac (a mark of enhancers and promoters) or H3K27me3 (the repressive mark deposited by EZH2 as a part of the PRC2 complex). We integrated publicly available H3K27ac/me3 ChIP-seq data generated from T24 cells [48, 49] and queried the number of genomic regions marked by each. Expectedly, most 100kb regions contained a peak for H3K27me3, with some regions marked by both H3K27me3 and H3K27ac, and a small proportion of regions marked by H3K27ac alone (Figure 4D). However, when we isolated the STAG2-bound regions only, the relative proportion of H3K27ac regions increased (Figure 4D, right) suggesting that STAG2 may specifically localize to regions marked with H3K27ac. Subsequent statistical testing confirmed that H3K27ac is more likely to occur at regions bound by STAG2 compared to regions that are not bound by STAG2, whereas H3K27me3 is more likely to be found in regions not bound by STAG2 (Figure 4E). To identify transcription factors likely to target genes in regions marked by STAG2-H3K27(ac/me3), we performed overrepresentation analysis using the MSigDB C3: Transcription Factor Targets gene sets. STAG2-H3K27ac co-bound genes are likely to be activated and are regulated by factors such as FOXD3 and BRF1 (Figure 4F, left), whereas STAG2-H3K27me3 co-bound genes are regulated by proteins such as ZNF436, which is reported to have repressor functions [50](Figure 4F, right).

### STAG2 directly binds DNA with co-regulators to modulate expression of invasion-related genes

We found that STAG2 is more likely to bind at A compartment chromatin, which is characteristically more transcriptionally active [22]. Therefore, we characterized the location of STAG2 binding peaks relative to the transcription start sites (TSSs) of coding genes. Relative to TSSs, STAG2 binds at a variety of regions, including promoters, introns, 3’ and 5’ UTRs, exons, as well as in distal intergenic regions (Figure 5A). For example, STAG2 binds at the promoter of the gene *ABCA1*, a cholesterol efflux pump known to promote cancer cell migration and invasion, which is also encoded in a region of significantly altered chromatin contact (Figure 5B, Figure 1B) [51, 52]. Next, we sought to identify possible STAG2 co-regulators. Utilizing the experimental technique of rapid immunoprecipitation with mass spectrometry of endogenous proteins (RIME), we detected proteins bound with STAG2 in the nucleus. The top binding partners included members of the cohesin complex, including STAG2, SMC3, SMC1A, RAD21, and cohesin accessory proteins PDS5A/B, WAPL, and NIBPL, demonstrating the effectiveness of the assay (Figure 5C, Table S9). To date, one previous publication has also investigated STAG2 binding partners through a mass spectrometry approach [53]. Therefore, we considered proteins which were identified in both experiments as high-confidence STAG2 binding partners (Figure 5D). Interestingly, the transcriptional co-repressor TRIM28 was identified in both datasets (Figure 5C-D, indicated in red) and has been previously found to colocalize with the cohesin complex in T-cells [25]. To query whether TRIM28 may co-regulate genes with STAG2 in patient samples, we performed regulatory transcriptional network (RTN) analysis on DEGs from the RP patient cohort. We found a significant enrichment for TRIM28 as a negative regulator of the DEGs that were identified by stratifying patients by STAG2 H-score, suggesting a co-regulatory relationship (Figure 5E). To confirm that STAG2 and TRIM28 co-localize to the same genomic regions in our model system, we performed ChIP for both proteins and a species-matched IgG negative control in T24 cells. We then amplified a genomic region within the ABCA1 promoter (indicated in Figure 5B) by quantitative real-time PCR (qRT-PCR). Results show that there is enrichment of binding of both STAG2 and TRIM28 over IgG negative control signal in both cell lines (Figure 5F), confirming their ability to target the *ABCA1* promoter. Finally, we sought to integrate our STAG2 binding data with our gene expression data to predict the gene regulatory function of STAG2. We integrated our STAG2 ChIP-seq dataset with differentially expressed genes after STAG2 knockdown via shRNA (Figure 5G, left) and total knockout (KO) via CRISPR-Cas9 (Figure 5G, right) utilizing the Cistrome-BETA plus analysis pipeline [54]. This pipeline assigns genes a regulatory potential score based on the magnitude of differential expression after STAG2 KD/KO combined with proximity to STAG2 binding sites. In both cases (STAG2 KD and STAG2 KO vs. control) the genes that were upregulated after STAG2 was lost were ranked slightly but significantly higher in regulatory potential score compared to genes which are not differentially expressed (Figure 5G). This result suggests that STAG2 may have a modest repressive effect on gene expression when present. Altogether these data support the hypothesis that STAG2 binds directly to DNA to regulate gene expression in concert with additional co-regulatory proteins.

**Figure 5.**
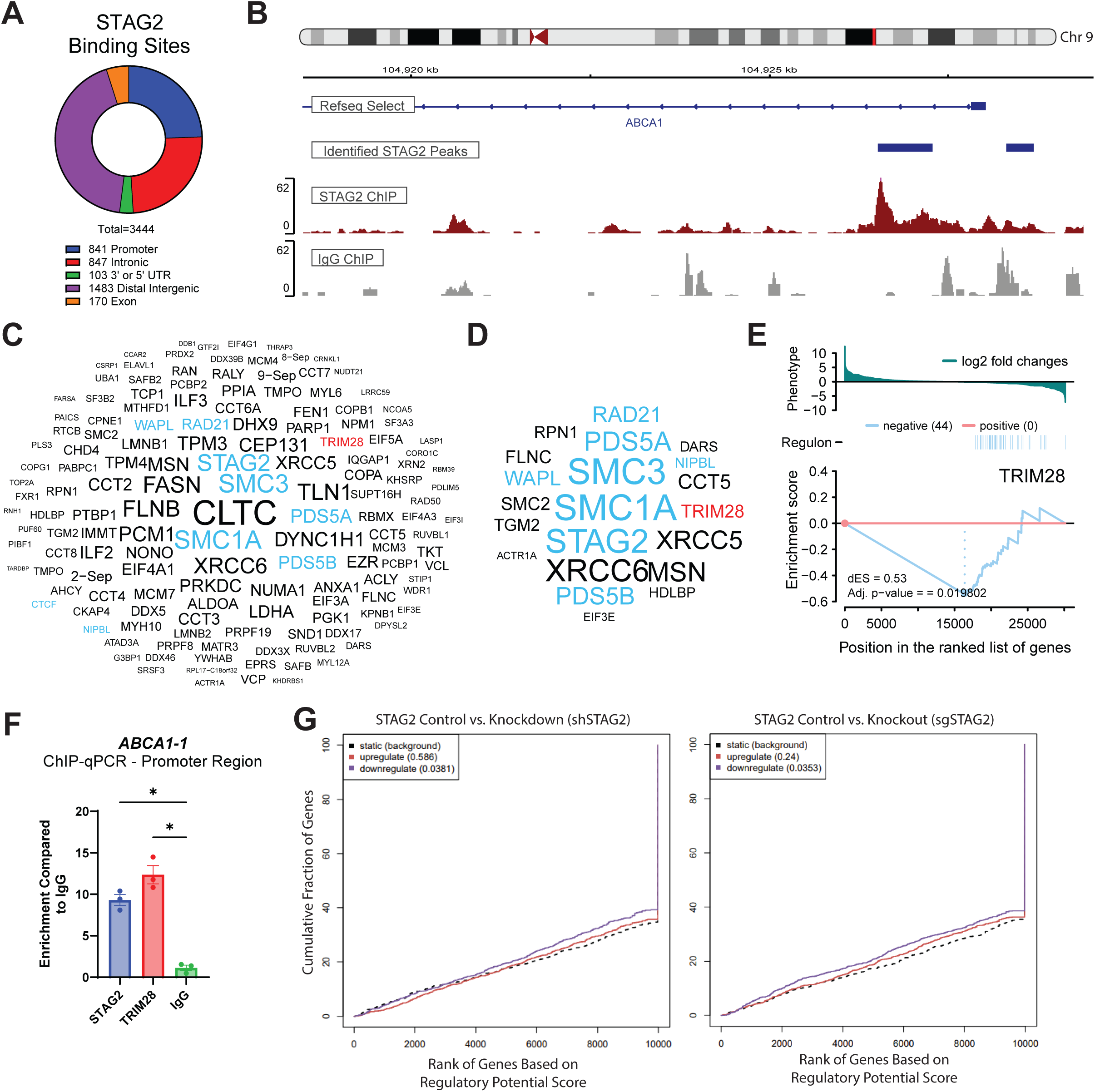
STAG2 directly binds DNA with co-regulators to modulate expression of invasion-related genes. A. Pie chart representing the different STAG2 binding sites identified by ChIP-sequencing in T24 cells. B. IgV visualization of STAG2 and IgG (negative control) binding at the candidate gene locus ABCA1 on chromosome 9. C. Word cloud depicting STAG2 co-bound proteins identified from RIME. Letter size represents percentage peptide spectral matches for each protein. D. Word cloud depicting STAG2 co-bound proteins identified from both our RIME and STAG2 RIME performed by [127]. Letter size represents percentage peptide spectral matches for each protein. For both word clouds: blue represents cohesin-complex members or known cohesin associated proteins; red represents protein of interest TRIM28. E. RTN analysis of differentially expressed genes (DEGs) from the RP patient cohort indicating predicted regulatory enrichment of transcriptional corepressor TRIM28 on genes enriched in STAG2 low. F. Enrichment for ABCA1 promoter amplification after ChIP for STAG2, TRIM28, or IgG in T24 MIBC cells. Data normalized to input signal per cell line and enrichment calculated against IgG signal per cell line. P-values assessed via one-way ANOVA, *p<0.05. G. Cistrome-BETA functional analysis outputs for STAG2 binding sites integrated with DEGs after shRNA-mediated STAG2 KD (left) or CRISPR-Cas9 mediated STAG2 KO (right). Red lines indicate genes upregulated after STAG2 KD/KO, blue represents genes downregulated after STAG2 KD/KO, black represents genes not differentially expressed (static). P-values for each group compared to background indicated in parentheses.

### STAG2 organizes chromatin and maintains histone acetylation to promote *ABCA1* expression and invasion

Hi-C, proteomics, ChIP-, and RNA-sequencing results suggested that STAG2 may regulate expression of genes through interactions with TRIM28, through maintenance of chromatin compartmentalization, and through control of chromatin contacts. To examine the mechanistic function of STAG2 at a more granular, single-gene level, we identified the candidate gene *ABCA1* as a model gene to investigate. *ABCA1* satisfied three major criteria: 1. *ABCA1* is STAG2-bound at the promoter and enhancer, as measured via ChIP-seq; 2. *ABCA1*is differentially expressed post-STAG2 KO; 3. ABCA1 has a known role in cancer cell invasion and migration as reported in the literature, thus providing a connection to the invasive phenotype observed in association with STAG2 in our models [51, 52].

From mass spectrometry analyses, we identified the transcriptional co-repressor TRIM28 as a possible STAG2 coregulatory protein. Therefore, we assessed co-occupancy of STAG2 and TRIM28 at the *ABCA1* promoter region via ChIP-qPCR. We found that TRIM28 promoter occupancy increases after STAG2 KO, suggesting a possible inverse functional role of TRIM28 and STAG2 on regulating *ABCA1* expression (Figure 6A). In line with the transcriptional corepressor function of TRIM28 [55, 56], increased TRIM28 promoter occupancy was associated with a significant downregulation of *ABCA1* mRNA and ABCA1 protein expression (Figure 6B-C). Given that TRIM28 can recruit histone deacetylases to remove acetylation at gene promoters [55, 57], we next assessed whether increased TRIM28 binding post-STAG2-KO was associated with changes in the histone landscape at *ABCA1*. Indeed, we observed downregulation of both H3K9 and H3K27 acetylation at the *ABCA1* promoter after STAG2 KO (Figure 6D). Given that acetylation marks are typically associated with active transcription [58, 59], the loss of H3K9ac and H3K27ac may mechanistically explain the downregulation of *ABCA1* after STAG2 KO.

**Figure 6.**
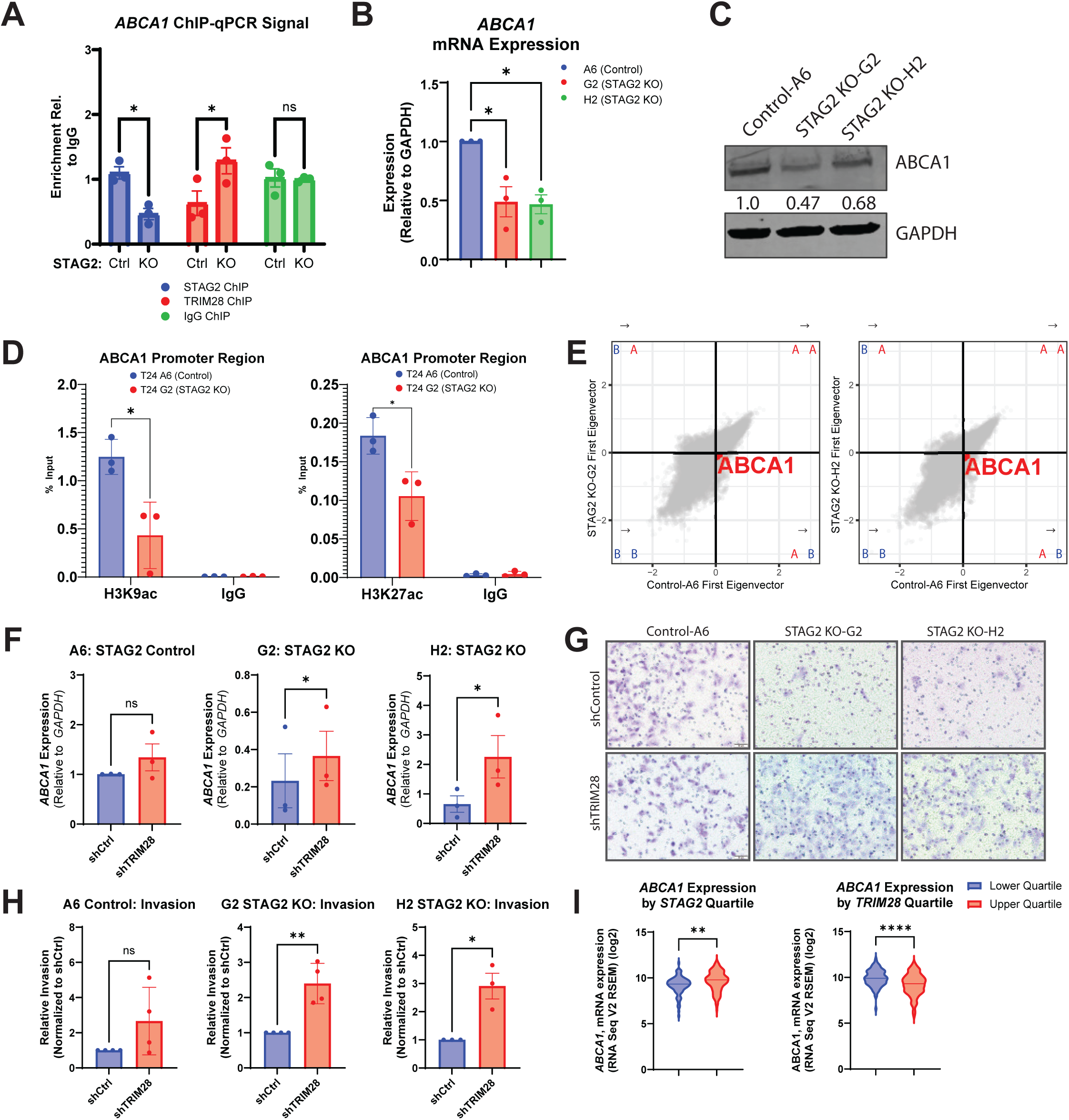
STAG2 organizes chromatin and maintains histone acetylation to promote ABCA1 expression and invasion. A. Enrichment for ABCA1 promoter amplification after ChIP for STAG2, TRIM28, or IgG in T24 Control (WT) and STAG2 KO cell lines. Data normalized to input signal per cell line and enrichment calculated against IgG signal per cell line. B. mRNA expression of ABCA1 relative to housekeeping gene GAPDH in T24 control and STAG2 KO cell lines as measured by qPCR. C. Western blot showing protein expression of ABCA1, GAPDH used as loading control and for quantification. D. Enrichment for ABCA1 promoter amplification after H3K9ac (left) and H3 K27ac (right) with IgG in T24 Control (WT) and STAG2 KO cell lines. Data normalized to input signal per cell line and enrichment calculated against IgG signal per cell line. E. Scatter plots depicting the first principal component score for each 10kb window across the genome, with positive score indicating compartment A, and negative score indicating compartment B for comparisons of T24 control A6 (STAG2 WT) versus T24 STAG2 KO G2 (left), and T24 control A6 versus T24 H2 KO (right). Percentage of 10kb-bins in each category (A in control and KO, B in control and KO, A in control and B in KO, and B in control A in KO) displayed. Genomic regions containing the ABCA1 locus are in red and reside in the A to B quadrant. F. mRNA expression of ABCA1 relative to housekeeping gene GAPDH in T24 control and STAG2 KO cell lines with concomitant shCtrl or shTRIM28 lentiviral constructs as measured by qPCR. G. Representative images of cells invaded through Matrigel-coated Transwell invasion chambers after 22 hours stained with crystal violet. H. Quantification of relative invasion for G. I. ABCA1 expression in TCGA Firehose Legacy BLCA cohort stratified by STAG2 (left) and TRIM28 (right) upper and lower quartile expression. *p<0.05, **p<0.01, ****p<0.0001.

We observed significant changes in chromatin features after STAG2 KO by Hi-C, and thus we next asked whether any chromatin changes occurred in regions containing the *ABCA1* gene. Indeed, the genomic region containing *ABCA1* experiences a compartment shift, going from A-compartment in the STAG2 control cells to B-compartment in both STAG2 KO cell lines (Figure 6E, *ABCA1* containing region marked with red dots). Altogether, we conclude that STAG2 expression is associated with A-compartment chromatin at *ABCA1*, which is further associated with a high degree of promoter H3K9 and H3K27 acetylation, which may contribute to elevated *ABCA1* expression when STAG2 is present.

To further examine a possible inverse regulatory function of STAG2 and TRIM28 on *ABCA1* expression, we used shRNA to knock down TRIM28 expression in all three of our T24 cell lines (Control A6, STAG2 KO G2, and STAG2 KO H2). We confirmed the knockdown of TRIM28 at both the protein and mRNA level in all three cell lines (Figure S5A-B). To measure changes in *ABCA1* expression, we performed qPCR for *ABCA1* in each cell line after TRIM28 KD. We found that the expression of *ABCA1* is rescued in the STAG2 KO cell lines after subsequent TRIM28 KD, suggesting a competitive interaction between the two proteins in regulating expression of *ABCA1* (Figure 6F). Further, the rescue of *ABCA1* expression post-TRIM28 KD was associated with a simultaneous restoration of invasive capability (Figures 6G-H). These results suggest that STAG2 and TRIM28 may inversely regulate *ABCA1* to control invasion in MIBC cells.

To validate that the inverse associations of STAG2 and TRIM28 with *ABCA1* hold true in clinical samples, we analyzed mRNA expression data from the TCGA BLCA cohort [60, 61].

Patients with the highest quartile of *STAG2* mRNA expression had significantly higher *ABCA1* expression, supporting a model in which STAG2 promotes expression of *ABCA1* (Figure 6I). Conversely, patients with the highest quartile of *TRIM28* mRNA expression had significantly lower *ABCA1* expression, supporting a repressive effect of TRIM28 on *ABCA1* (Figure 6I). Both results support a model in which STAG2 and TRIM28 may inversely regulate expression of *ABCA1* in MIBC.

### STAG2 localizes to a subset of PRC2 target genes to augment gene repression

We found that STAG2 and the PRC2-deposited repressive mark, H3K27me3, are frequently found within the same 100-kb genomic region (Figure 4E), suggesting that STAG2 and the PRC2 complex may co-regulate expression of certain genes. To identify a list of candidate co-regulatory genes, we intersected the EZH2 transcription factor target gene list from ENCODE [62, 63] with STAG2-repressed DEGs from T24 RNA-seq. We identified five candidate co-repressed genes: *ASAH1, CHRM3, LAMB2, RAB20,* and *SPOCK3,* which each had a positive H3K27me3 peak score within the respective genomic regions according to H3K27me3 ChIP-seq (Figure 7A). *SPOCK3* was the most differentially expressed gene after STAG2 KO (Figure 7B). Further, the *SPOCK3* gene experiences a B to A compartment shift after STAG2 KO (Figure 7C), has several STAG2 binding peaks (Figure 7D), and gains chromatin loops after STAG2 KO (Figure 7E). In prostate cancer, SPOCK3 expression minimizes invasion and migration is associated with better clinical outcomes [64]. Thus, we chose to investigate *SPOCK3* as a candidate PRC2-STAG2 co-repressed gene.

**Figure 7.**
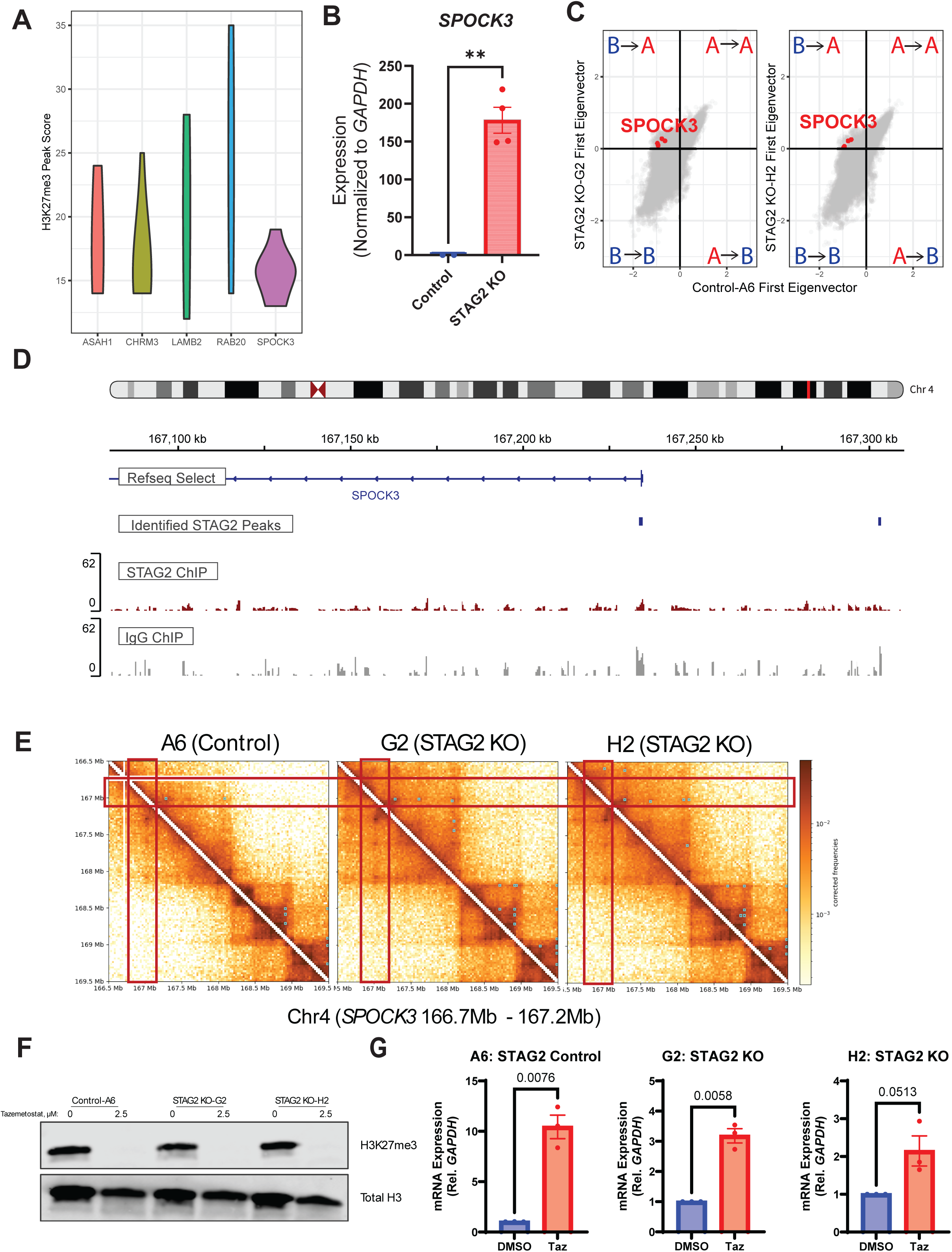
**STAG2 co-regulates PRC2 target genes and loss of STAG2 augments the effect of EZH2i on gene expression**. A. H3K27me3 peak score as assessed by ChIP-seq in regions containing five candidate STAG2-PRC2 target genes. B. *SPOCK3* mRNA expression in Reads Per Kilobase of transcript per Million mapped reads (RPKM) between control and STAG2 KO cells; **p<0.01, assessed by two tailed t-test. C. Scatter plots depicting the first principal component score for each 10kb window across the genome, with positive score indicating compartment A, and negative score indicating compartment B for comparisons of T24 control A6 (STAG2 WT) versus T24 STAG2 KO G2 (left), and T24 control A6 versus T24 H2 KO (right). Genomic regions containing the SPOCK3 locus (n=4) are in red and reside in the B to A quadrant. D. IgV visualization of STAG2 and IgG (negative control) signal at the candidate gene locus *SPOCK3* on chromosome 4. E. Contact frequency heatmaps for a 3 Mb region of chromosome 4; red boxes outline the region encoding SPOCK3. Teal dots represent focal enrichment, indicating strong chromatin looping events at the pairs of genomic regions indicated. F. Western blot showing protein expression of H3K27me3 in isolated histone fractions after 2.5 µM tazemetostat treatment; total histone H3 used as loading control. G. mRNA expression of *SPOCK3* relative to housekeeping gene GAPDH in T24 control and STAG2 KO cell lines after treatment with 2.5 µM tazemetostat measured by qPCR.

To test the effects of altered PRC2 activity on *SPOCK3*, we treated T24 cells with the EZH2 inhibitor tazemetostat. After 72-hours of treatment with 2.5 tazemetostat, H3K27me3 was absent from cells as evidenced by western blot (Figure 7F), showing on-target effects of the drug in PRC2 inhibition. After treatment, we assessed *SPOCK3* expression by qPCR. We found that STAG2 KO alone led to significant upregulation of *SPOCK3,* and that this effect was augmented even further by tazemetostat treatment (Figure 7G). Altogether, these results indicate that *SPOCK3* is co-repressed by STAG2 and PRC2. Given the previously established role for SPOCK3 in repressing invasion, *SPOCK3* upregulation after STAG2 KO may contribute to the diminished invasive potential that we observe *in vitro*.

## DISCUSSION

Understanding the complex role of STAG2 in cancer is imperative, given the high frequency of alterations in STAG2 across many cancer types [4, 5, 8, 65]. Conversely to other cancer types, expression of STAG2 is demonstrably pro-oncogenic in MIBC. Our findings suggest that this context-specific role may be due to the function of STAG2 in mediating chromatin organization and epigenetic characteristics that are specific to individual cell types. When comparing bladder tumor to adjacent normal bladder tissues, the cohesin chromatin regulation pathway was one of the top differentially enriched pathways. We also identified tumor-intrinsic differences dictated by STAG2 expression level, with STAG2 associated with transcriptional enrichment of pro-oncogenic signaling pathways. These data predicate a role for cohesin and STAG2 in bladder cancer initiation and progression and provide significant rationale for investigating the specific role of STAG2 in MIBC. STAG2 assists with chromatin organization as part of the cohesin complex in both malignant and non-malignant mammalian cells [3–6, 19–21]. However, the contributions of STAG2 to 3D chromatin architecture, including chromatin contact frequency, compartmentalization, looping, and associations of these characteristics with gene expression have not been investigated in MIBC.

Here, we find that STAG2 loss results in widespread changes in contact frequency between pairs of genomic regions. The number of pairs gaining contact was nearly double the number of pairs losing contact with each other after STAG2 KO (1,011 pairs gained vs. 566 pairs lost). This suggests that STAG2 frequently acts as a genomic insulator, preventing aberrant contact between pairs of genomic regions, and to a lesser degree can also maintain proximity of genomic pairs. When investigating the distance between pairs of genomic regions that gained or lost contact with each other, we found that genomic regions that gained contact were significantly further apart, whereas lost contacts tended to be closer together in the linear genome. STAG2 KO resulted in loss of shorter, weaker chromatin loops and a gain of longer-ranging, stronger chromatin loops. This is supported by previous work showing that STAG2 is responsible for short-range genomic contacts, whereas STAG1 is responsible for longer-range genomic contacts [20, 44]. STAG2-cohesin binds more transiently to DNA compared to STAG1-cohesin [66]. The difference in residence time on DNA between STAG1- and STAG2-cohesin may explain the propensity for STAG2 in mediating shorter ranged contacts, as a shorter residence time would result in a shorter time window to extrude DNA into chromatin loops, resulting in smaller loop formation [67]. Our results suggest a remodeling of chromatin contact and looping that leads to longer-ranged, stronger interactions which may be dominated by STAG1-cohesin after STAG2 KO.

Chromatin compartmentalization segregates the genome into regions of open and active versus closed and inactive chromatin [22]. After STAG2 KO, a larger proportion of regions shifted from the repressed B compartment to the active A compartment, though shifts were observed in both directions. This suggests that STAG2 may repress gene expression in many cases by maintaining B-compartment chromatin, while promoting active gene expression in other regions by maintaining A-compartment chromatin. When investigating the direct effects of altered chromatin compartmentalization, we find a strong and direction-specific correlation between the magnitude of compartment shift and the magnitude of gene expression change. We also identified a significant association between regions with altered contact frequency and differential expression of the genes encoded in these respective regions. This demonstrates a direct contribution of STAG2-mediated chromatin contact frequency and chromatin compartmentalization in regulating level of gene expression.

STAG2 localization varies based on cell type, which may partially explain how STAG2 regulates cell type-specific gene expression programs [18, 19]. However, the mechanisms underlying this specificity in binding remain unclear. Previous work demonstrates that STAG2 and cohesin may colocalize with epigenetic regulators such as TRIM28 and Polycomb complexes, which regulate expression through altering the abundance of specific histone post-translational modifications [19]. In our model system, we found that sites bound by STAG2 have proportionally more H3K27ac than non-STAG2 sites. The H3K27ac mark defines active chromatin regions [62], implying that STAG2 preferentially binds at active chromatin. This suggests that STAG2 localization may be dictated by histone post-translational modifications or “marks,” which vary widely between cell types. This may explain the context-specific effects of STAG2 loss in different cancer types, which should be further investigated via comparisons of STAG2 localization across different cancer contexts.

STAG2 binding was also significantly associated with compartment state, with STAG2 more likely to bind in regions designated as A-compartment. This further demonstrates a propensity for STAG2 binding at active chromatin. Interestingly, we found that after STAG2 KO the magnitude of B to A shifts is greater than A to B. This may be because STAG1 can compensate for STAG2 loss in active chromatin regions, but cannot compensate in less active chromatin regions, causing a greater proportion of changes in B-compartment regions after STAG2 KO. Interestingly, we found that STAG2 maintains B-like chromatin most frequently at distal enhancer regions. This supports a model in which STAG2 localizes frequently at A-compartment chromatin while simultaneously maintaining B-compartment chromatin at distal enhancers. This phenomenon may be explained by a specificity for STAG2 binding at specific histone post-translational modifications. Additionally, increased contact with distal enhancer regions after STAG2 KO was frequently associated with changes in gene expression. This further implicates a role for STAG2 in preventing aberrant enhancer activity through maintenance of B-compartment chromatin and minimizing gene-enhancer contacts to restrict gene expression. As enhancer reprogramming is a prominent feature in cancer [68], we propose that STAG2-mediated enhancer reprogramming may drive tumor-promoting gene expression programs, contributing to worse outcomes for patients with high STAG2 expression.

We found significant alterations in chromatin features after STAG2 KO, which we hypothesized would have consequences on expression of STAG2-regulated genes. At individual gene loci, we found that STAG2 has divergent mechanisms depending on the directionality of gene regulation; that is, whether the expression is promoted by STAG2, or the expression is repressed by STAG2. Loss of STAG2 at the STAG2-promoted gene *ABCA1* is associated with loss of H3K9/27ac, increased TRIM28 occupancy, and gene downregulation. These changes in histone acetylation may contribute to shifts in compartmentalization, as we also found that *ABCA1* experiences an A to B compartment shift after STAG2 KO. Whether these changes in acetylation are due to increased TRIM28 occupancy leading to HDAC recruitment, or due to altered HDAC or HAT expression because of STAG2 loss, remains to be investigated.

As a secondary mechanism of gene regulation, we identified a set of STAG2 repressed genes that are reported to be targeted by PRC2 and have high levels of H3K27me3 in T24 cells. We investigated the STAG2-PRC2 target gene *SPOCK3* and found that loss of STAG2 led to aberrant loop formation surrounding the *SPOCK3* gene locus and resulted in *SPOCK3* upregulation. This was further augmented by treatment with the EZH2 inhibitor tazemetostat, suggesting cooperation between STAG2 and PRC2 in gene repression. This is in line with evidence for Polycomb and cohesin complex cooperation in other models [5, 19, 25–27].

Given our results showing an association between altered histone modifications and expression of STAG2-regulated genes, we speculate that inhibition of these marks (via histone acetyltransferase inhibitors and/or EZH2 inhibitors) strategies to diminish the activity and pro-oncogenic function of STAG2. Altogether, this work contributes a novel understanding of how STAG2 maintains expression of pro-tumor gene expression programs through maintaining specific chromatin characteristics at genes and regulatory elements, restricting aberrant contact with distal enhancers, and collaboration with epigenetic enzymes to regulate histone post-translational modifications. These findings may underlie the context-specific function of STAG2 across cancer types and lay the groundwork for investigating epigenetic inhibitors as STAG2-specific therapeutic approaches.

## CONCLUSIONS

STAG2 maintains chromatin compartmentalization states and restricts interactions between genes and regulatory elements to promote oncogenic gene expression programs. At specific genes, STAG2 interacts with epigenetic regulators such as TRIM28 and the PRC2 complex to exert divergent gene regulatory functions through modulation of histone modifications in combination with regulation of chromatin state. These findings identify novel mechanisms of STAG2-mediated gene regulation that occur in MIBC that may help explain the divergent functions of STAG2 across cancer types and provide rationale for investigation of epigenetic inhibitors as STAG2-specific precision medicine approaches.

## ACKNOWLEDGEMENTS

Ethical Approval and Consent to Participate

All tissue samples were obtained with written informed consent from each patient to allow the use of his/her/their samples and data in research.

**Acknowledgments**

This work was supported by National Cancer Institute (NCI) grant P30CA016056 involving the use of Roswell Park Comprehensive Cancer Center’s Shared Resources.

## Author contributions

Conceptualization: SRA, DGT, GG, AW; Methodology: SRA, AS, SLV, PS, MPV, GG, AW. Formal analysis: SRA, JZ, XL, ECG, BD, JJ, SLV, PS, MPV; Investigation: SRA, BD, AS, SLV, MPV; Writing – original draft: SRA, AW; Writing – review & editing: SRA, AW; Visualization: SRA, JZ, XL, ECG, BD, JJ, TL; Supervision: AW; Funding acquisition: SRA, AW. All authors have read and approved the final version of the manuscript and agree to be accountable for all aspects of the work.

## Data availability statement

The genomics data generated for this study are publicly available in the Gene Expression Omnibus (GEO) database (https://www.ncbi.nlm.nih.gov/geo/) under accession numbers GSE329907 (Hi-C) and GSE334379 (T24 RNA-sequencing and T24 STAG2 ChIP-sequencing).

## SUPPLEMENTARY FIGURE AND TABLE LEGENDS

### Supplementary Figure Legends

**Figure S1.** A. IPA canonical pathway enrichment analysis of differentially expressed genes between bladder tumor (n = 66) and adjacent non-tumor/normal (n = 15) samples from the RP cohort. Pathways are ranked by -log10(P-value). B. H-score (histo-score, representative of STAG2 protein expression level) for STAG2-low and STAG2 high adjacent non-tumor (normal, N) in blue, and bladder tumor (T) samples in red. P-value assessed by t-test, ** p<0.01, **** p<0.0001. C. Correlation between STAG2 protein expression (H-score) and log2-transformed STAG2 mRNA expression in tumor and adjacent non-tumor/normal samples from the RP cohort. Pearson correlation coefficiency (r) and corresponding p-value are shown. D. Kaplan-Meier overall survival (left) and disease-free survival (right) curves for the TCGA BLCA cohort stratified based on median STAG2 mRNA expression. E. Kaplan-Meier survival curve for the Roswell Park patient cohort based on STAG2 signature score, comparisons made between the top 25% signature scored patients (n = 11) versus the bottom 75% (n = 30) signature scored patients. Survival measured in months. P-value determined by log rank test.

**Figure S2.** A. Left: Representative images and right: quantification of cells (for STAG2 Control A6 (STAG2 WT), STAG2 KO G2, and STAG2 KO H2) invading through Matrigel-coated Transwell membranes, stained with crystal violet. Right: quantification of invasion across three experimental replicates relative to control A6 cell line. B. Whole-genome contact matrices and C. single chromosome (chromosome 1) contact matrices for STAG2 Control A6 (STAG2 WT), STAG2 KO G2, and STAG2 KO H2 cell lines. Comparisons made using one-way ANOVA; ***p<0.001. D. Total distribution of contact changes after STAG2 KO for pairs of genomic regions across the entire genome. E. Distribution of contact frequency changes by chromosome after STAG2 KO. Red: regions of increased contact frequency post-STAG2 KO; blue, regions of decreased contact frequency post-STAG2 KO. Only regions of statistically significant changes in contact shown for all plots.

**Figure S3.** A. Proportions of regions for each compartment state by chromosome. B. Proportions of STAG2-bound regions for each compartment state by chromosome. C. Proportions of STAG2-bound shifted regions by chromosome. D. Scatter plot of average shift in compartment and Log2 Fold Change for STAG2-bound regions encoding genes experiencing compartment shifts, each dot representative of a single gene. AA: A remaining A compartment; AB: A to B compartment switch; BA: B to A compartment switch; BB: B remaining B compartment.

**Figure S4.** A. Proportion of regions for each compartment state by cis-regulatory element type (CRE type). B. Proportion of only compartment shifted regions by CRE type. C. Proportion of STAG2 bound regions by CRE type. D. Proportion of STAG2-bound region compartment states by CRE type. E. Proportion of STAG2-bound compartment shifted regions by CRE type. AA: A remaining A compartment; AB: A to B compartment switch; BA: B to A compartment switch; BB: B remaining B compartment.

**Figure S5.** A. Western blot of T24 control and STAG2 KO cell lines with either shCtrl or shTRIM28, probed for TRIM28 and GAPDH as loading control. B. TRIM28 mRNA expression relative to GAPDH in T24 A6 Control (left), STAG2 KO G2 (middle) and STAG2 KO H2 (right) shCtrl and shTRIM28 cell lines. Statistics evaluated by two-tailed t-test, *p<0.05; **p<0.01.

### Supplementary Table Legends

**Table S1. RP DEGs, Tumor vs. Adjacent Non-Tumor/Normal Tissues.** All genes that are differentially expressed between bladder tumor (n = 66) versus normal (n = 15) tissues within the Roswell Park Comprehensive Cancer Center patient cohort. Only genes with an absolute log2FoldChange > 4 and padj < 0.05 are shown.

**Table S2. RP DEGs, STAG2 H-Score High vs. Low.** All genes that are differentially expressed between STAG2-high bladder tumors (n = 17) versus STAG2-low bladder tumors (n = 38) within the Roswell Park Comprehensive Cancer Center patient cohort.

**Table S3. TCGA DEGs, STAG2 mRNA High vs. Low.** All genes that are differentially expressed between *STAG2*-high bladder tumors (n = 204) versus *STAG2*-low bladder tumors (n = 204) from the TCGA BLCA bladder cancer cohort. Samples were classified based on median expression of *STAG2* mRNA.

**Table S4. T24 DEGs, STAG2 WT vs. STAG2 KO.** All genes that are differentially expressed between T24 control-A6 (STAG2 WT) cells compared to T24 STAG2 KO-G2 and STAG2 KO-H2 cells.

**Table S5. Hi-C Differential Contacts.** List of all 100-kb genomic region pairs with altered contact for STAG2 KO clone G2 vs control A6 (sheet 1) and STAG2 KO clone H2 vs Control A6 (sheet 2).

**Table S6. Dots Identified by Hi-C.** Enriched foci or “dots” identified in T24 Control-A6 cells (sheet 1), STAG2 KO-G2 cells (sheet 2), and STAG2 KO-H2 cells (sheet 3).

**Table S7. Insulator Region Quantifications.** Insulator/boundary quantifications for 100kb regions in T24 Control-A6 cells (sheet 1), STAG2 KO-G2 cells (sheet 2), and STAG2 KO-H2 cells (sheet 3).

**Table S8. 1st Eigenvector Scores for Chromatin Compartment Analysis.** First eigenvectors values for 100 kb regions in in T24 Control-A6 cells (sheet 1), STAG2 KO-G2 cells (sheet 2), and STAG2 KO-H2 cells (sheet 3). Positive eigenvector represents A-compartment chromatin, negative first eigenvector represents B-compartment chromatin.

**Table S9. STAG2 Nuclear Interacting Proteins by Mass Spectrometry.** Nuclear STAG2 interacting proteins as identified from immunoprecipitation followed by mass spectrometry.

