## Supplementary figures and images for "STAG2 Maintains Chromatin Compartmentalization and Represses Regulatory Element Contact to Promote Oncogenic Signaling in Muscle Invasive Bladder Cancer"

### Supplemental Figures

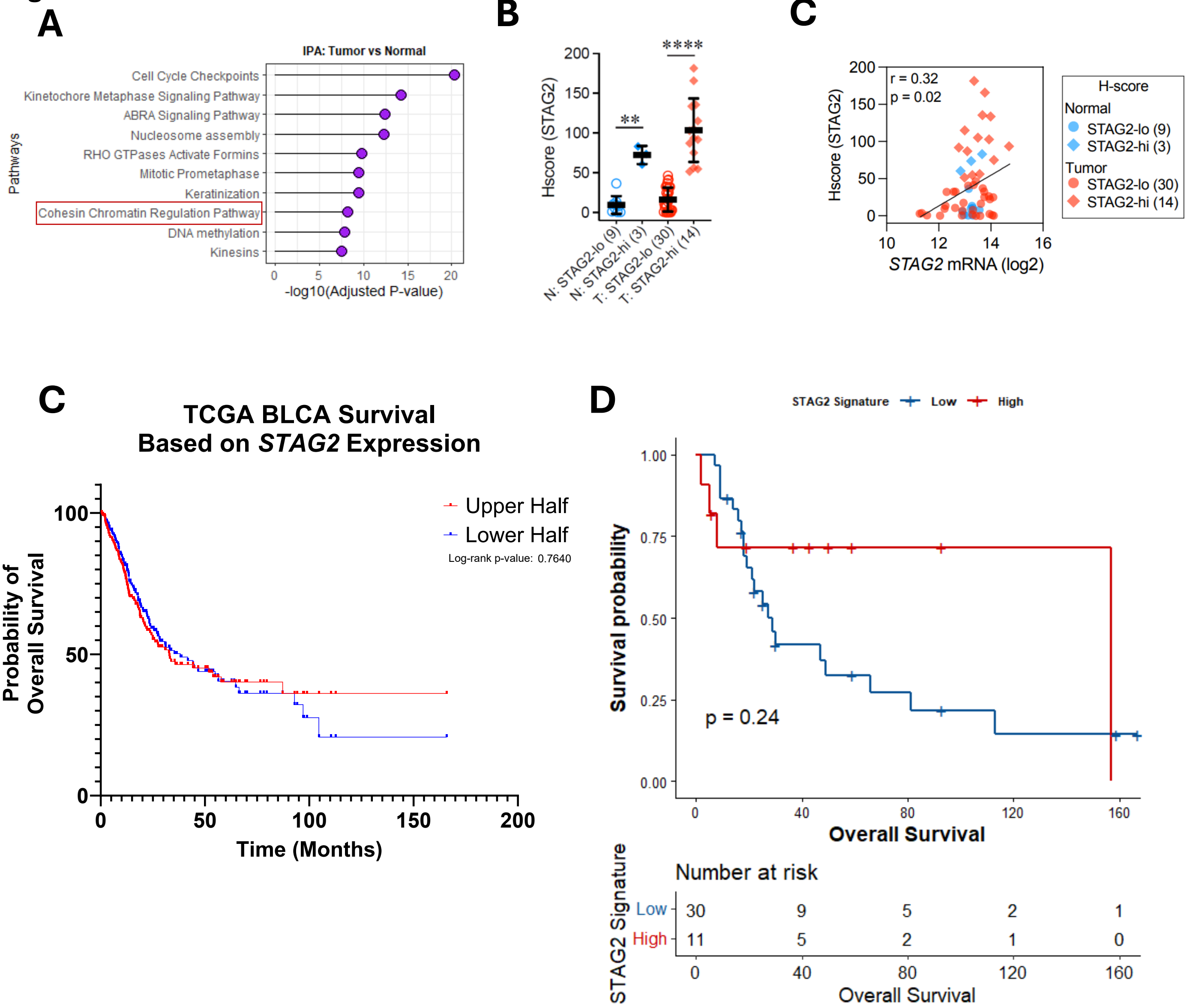

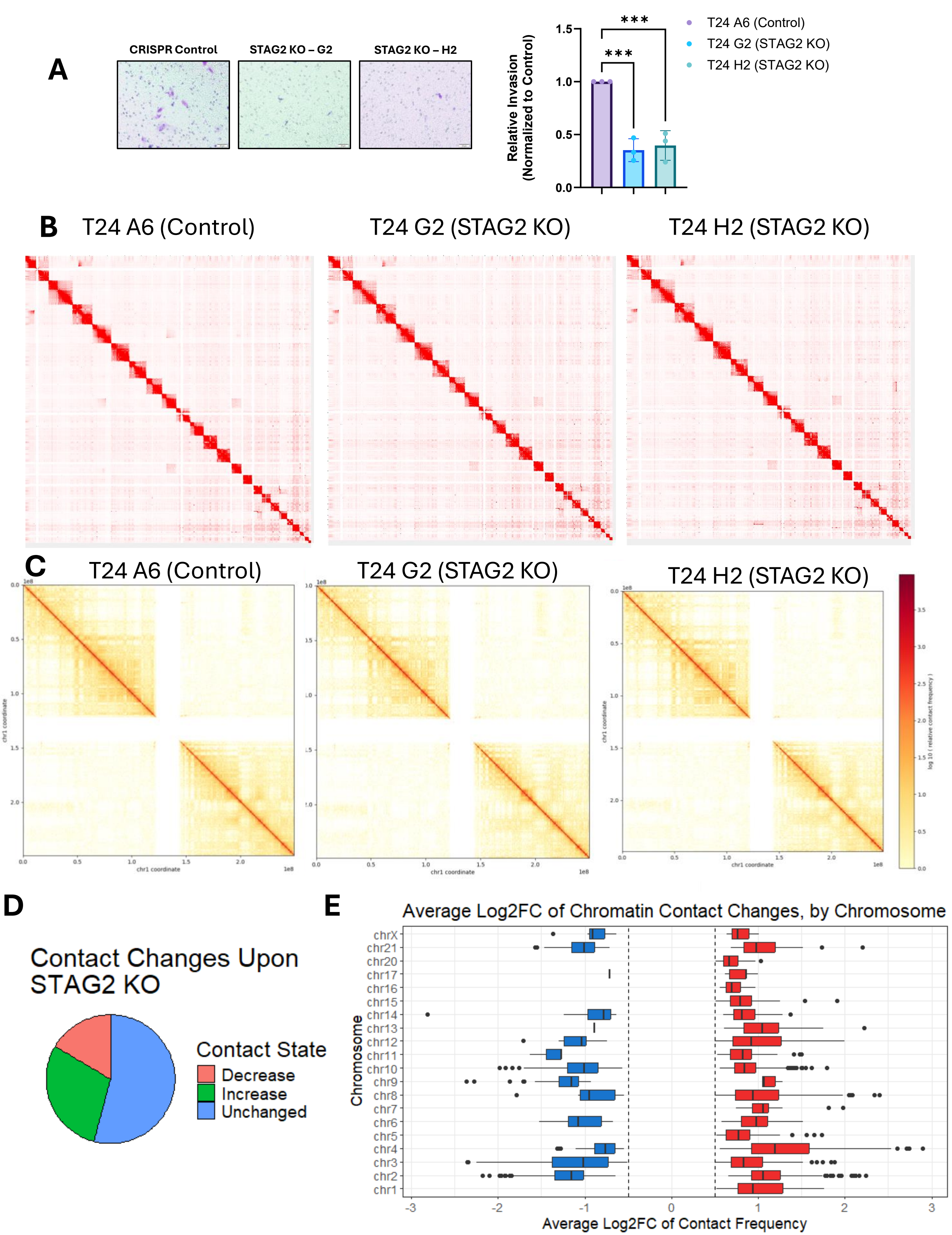

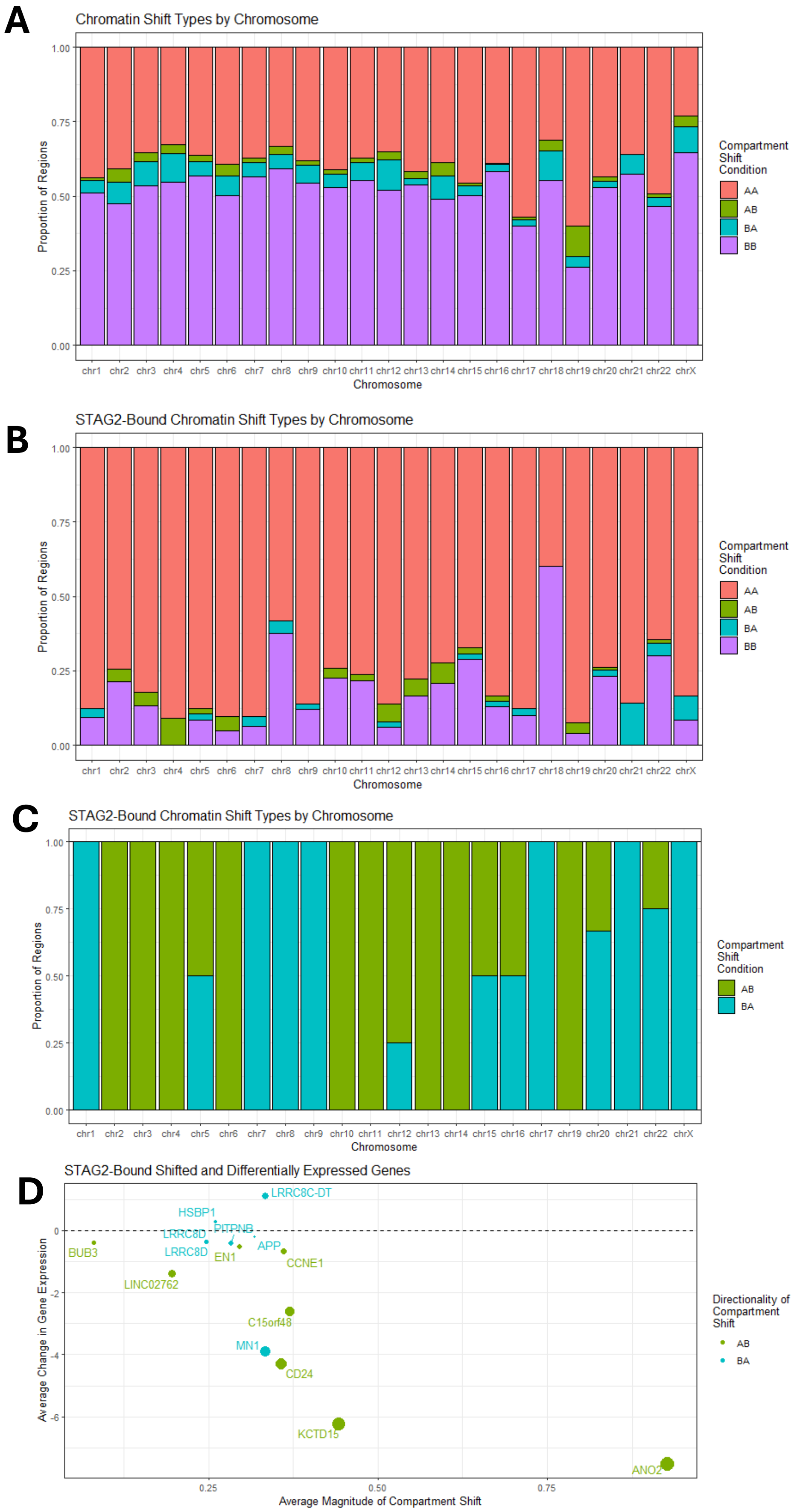

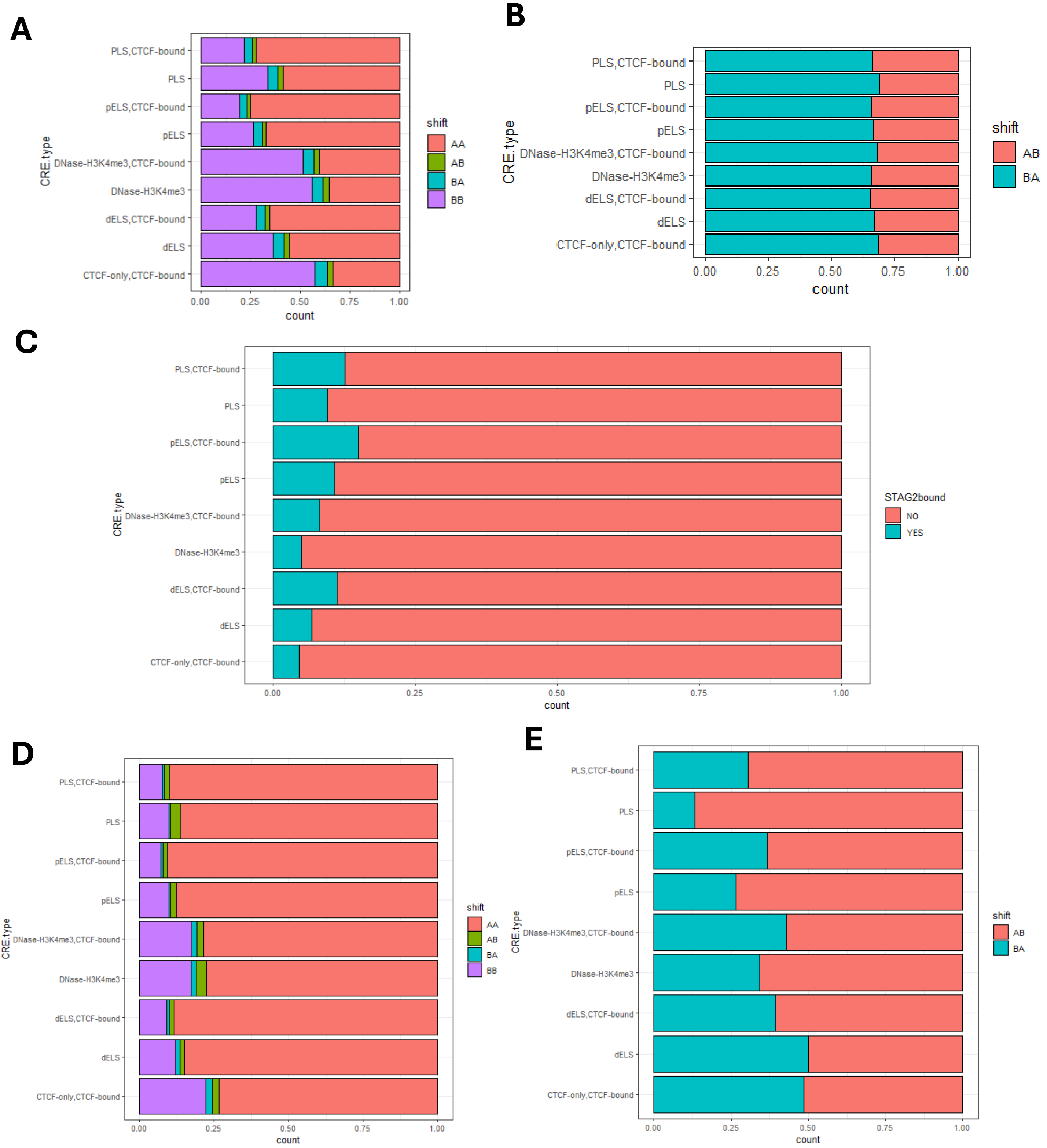

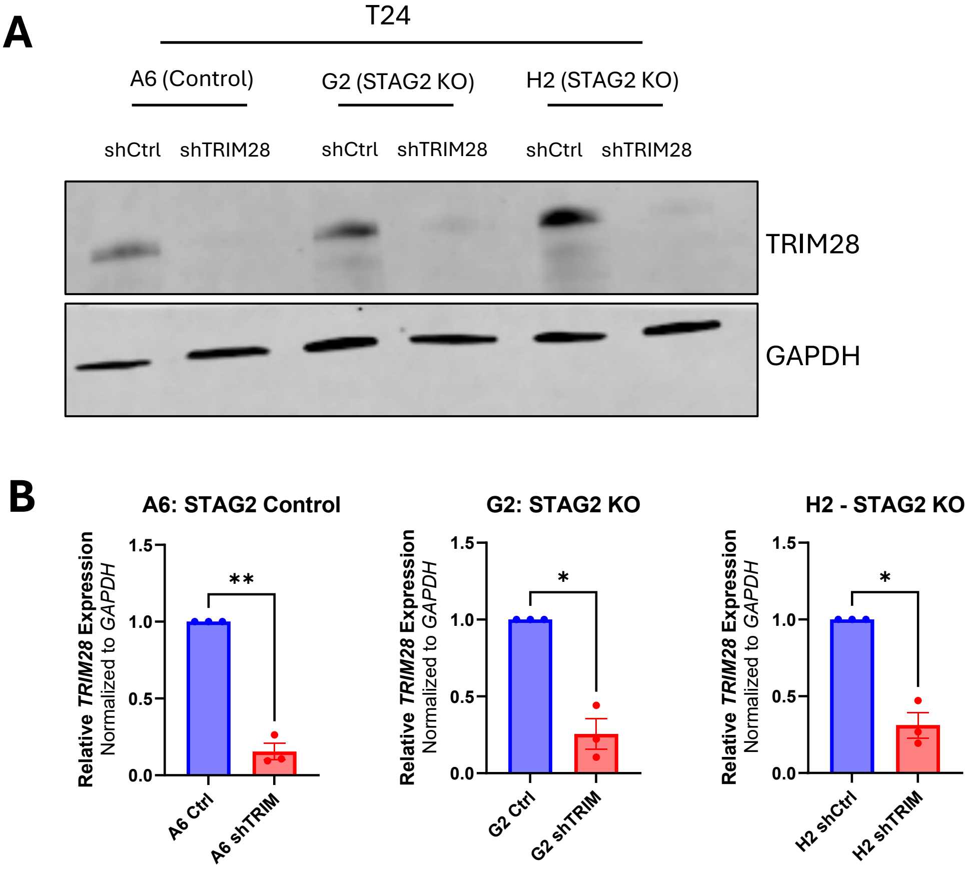
